# Characterizing Subcortical Structural Heterogeneity in Autism

**DOI:** 10.1101/2023.08.28.554882

**Authors:** David N. MacDonald, Saashi A. Bedford, Emily Olafson, Min Tae M. Park, Gabriel A. Devenyi, Stephanie Tullo, Raihaan Patel, Evdokia Anagnostou, Simon Baron-Cohen, Edward T. Bullmore, Lindsay R. Chura, Michael C. Craig, Christine Ecker, Dorothea L. Floris, Rosemary J. Holt, Rhoshel Lenroot, Jason P. Lerch, Michael V. Lombardo, Declan G. M. Murphy, Armin Raznahan, Amber N. V. Ruigrok, Elizabeth Smith, Russell T. Shinohara, Michael D. Spencer, John Suckling, Margot J. Taylor, Audrey Thurm, MRC AIMS Consortium, Meng-Chuan Lai, M. Mallar Chakravarty

## Abstract

Autism presents with significant phenotypic and neuroanatomical heterogeneity, and neuroimaging studies of the thalamus, globus pallidus and striatum in autism have produced inconsistent and contradictory results. These structures are critical mediators of functions known to be atypical in autism, including sensory gating and motor function. We examined both volumetric and fine-grained localized shape differences in autism using a large (*n*=3145, 1045-1318 after strict quality control), cross-sectional dataset of T1-weighted structural MRI scans from 32 sites, including both males and females (assigned-at-birth). We investigated three potentially important sources of neuroanatomical heterogeneity: sex, age, and intelligence quotient (IQ), using a meta-analytic technique after strict quality control to minimize non-biological sources of variation. We observed no volumetric differences in the thalamus, globus pallidus, or striatum in autism. Rather, we identified a variety of localized shape differences in all three structures. Including age, but not sex or IQ, in the statistical model improved the fit for both the pallidum and striatum, but not for the thalamus. Age-centered shape analysis indicated a variety of age-dependent regional differences. Overall, our findings help confirm that the neurodevelopment of the striatum, globus pallidus and thalamus are atypical in autism, in a subtle location-dependent manner that is not reflected in overall structure volumes, and that is highly non-uniform across the lifespan.

## Introduction

While the exact etiology has remained elusive, autism is a neurodevelopmental condition that significant evidence suggests is associated with neuroanatomical and functional alterations. Nevertheless, studying the neurobiology of autism is difficult due to the great deal of heterogeneity in autism trait profiles and co-occurring conditions; there are also no reliable biomarkers that indicate the presence of autism. Autism presentation varies greatly over the lifespan, by sex (throughout this manuscript, the term “sex” refers to sex assigned at birth). While autism presentation is three to four times more prevalent in males than in females, females with autism are more likely to be severely affected [1–3]. Even commonly co-occurring conditions present in different combinations in different individuals, and some, such as intellectual disability, can greatly affect the presentation [1]. This behavioural heterogeneity is consistent with the lack of agreement in the literature about neurobiological alterations in autism. In this study, as a followup to studies by our group focused on the cortex [4, 5], we attempt to address this by examining autism at the subcortical level. We account for three potential sources of neurobiological heterogeneity: sex, age, and full-scale intelligence quotient (FIQ), since variation along each of these dimensions have been associated with variation in both behavioural characteristics of autism and neurodevelopment [6–8].

Given the central role of the thalamus, striatum, and globus pallidus in behaviours known to be affected in autism, and the evidence of structural and functional differences in autism [9–11], they continue to remain understudied in comparison to the cortex. There are reports that, compared to typically developing individuals, the volume of the thalamus in individuals with autism is larger [12], smaller [13–15], and not significantly different [9, 16–19]. Similarly, the striatum has been reported to be larger [16, 20, 21], smaller [14, 22], and not significantly different [9, 13, 17–19]. The findings vary as well for the globus pallidus [9, 13, 14, 16–19, 22]. Noting a similar phenomenon at the cortical level, Bedford et al. [4] proposed three potential sources of this lack of concordance: 1) heterogeneity within the autism samples that had not been parsed, 2) issues with data quality such as motion artifact, and 3) the use of gross volumetric measures that may obscure subtle, localized differences in structure area or shape.

The overarching goal of this project is to resolve the disagreements about the nature of subcortical anatomical differences in autism, using spatially-sensitive techniques that account for variation due to age, sex, and IQ, in a rigorously quality-controlled, large, highly-powered multisite dataset. In doing so, we hope to parse some of the heterogeneity that may be interfering with finding clear results, and to reduce the effects of spurious, inter-site differences.

## Methods

### Sample

Analysis was performed on a large, multisite dataset of T1-weighted MRI scans of the head, which has previously been characterized by our group at the cortical level [4, 5]. The demographic composition of this dataset is detailed in Tables 1 and 2, but in brief, of the 3145 participants, ranging in age from 2-65 years, 1415 have a diagnosis of autism (1165 male/250 female) and 1730 are considered typically developing (TD: 1172 male/558 female). The dataset comprises both releases of the multi-site ABIDE I/II dataset [23, 24], as well as data from the National Institute of Mental Health (USA), the Hospital for Sick Children (Canada), the Cambridge Family Study of Autism (UK), and the UK Medical Research Council Autism Imaging Multicentre Study (UK), for a total of 32 sites.

**Table 1:**
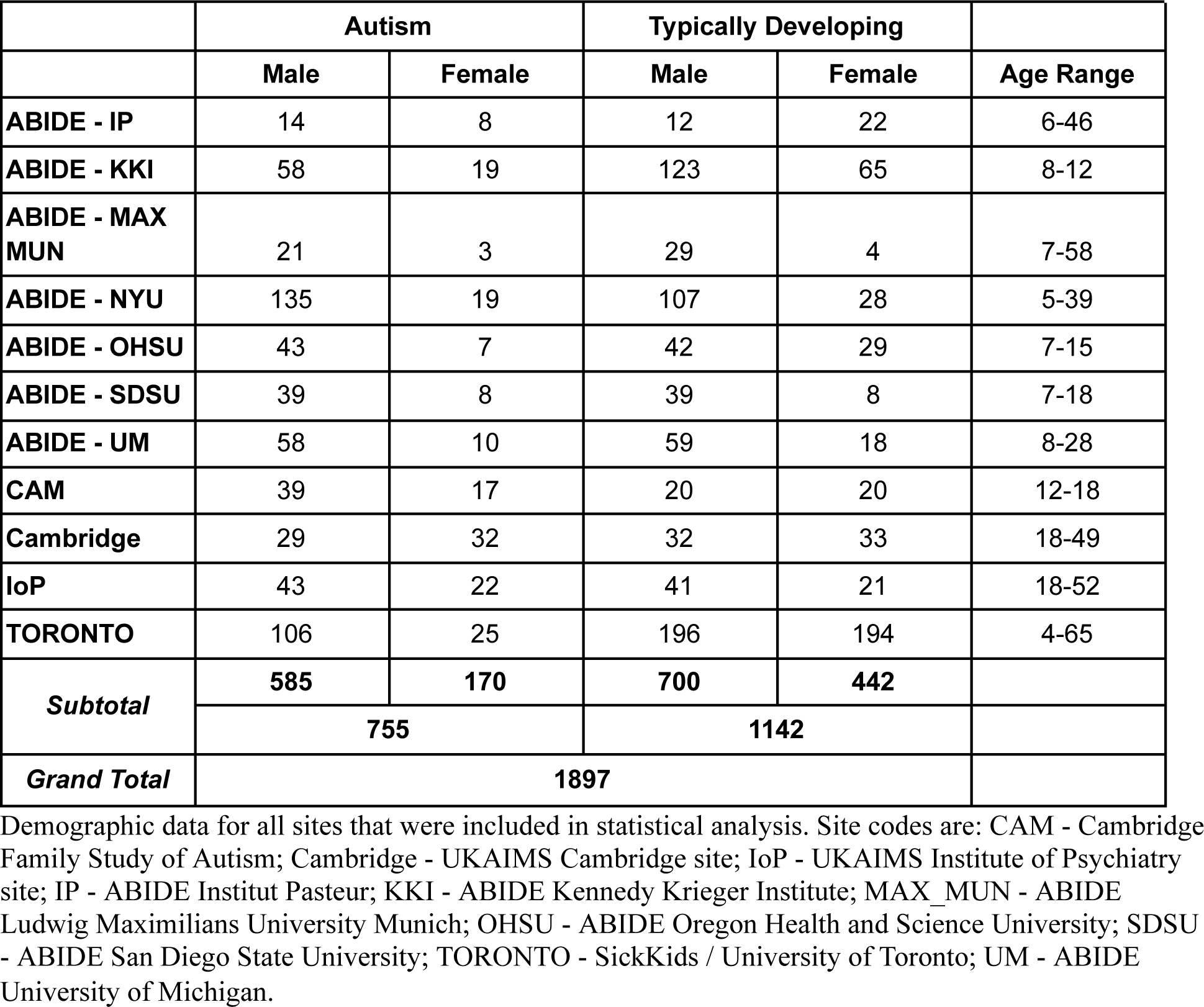
Demographics (Included Sites)

**Table 2:** Demographics (Excluded Sites)

|  | Autism |  | Typically Developing |  |  |
| --- | --- | --- | --- | --- | --- |
|  | Male | Female | Male | Female | Age Range |
| ABIDE - BNI | 29 | 0 | 29 | 0 | 18-64 |
| ABIDE - CALTECH | 15 | 4 | 15 | 4 | 17-56 |
| ABIDE - CMU | 11 | 3 | 10 | 3 | 19-40 |
| ABIDE - EMC | 22 | 5 | 22 | 5 | 6-10 |
| ABIDE - ETH | 13 | 0 | 24 | 0 | 13-30 |
| ABIDE - GU | 43 | 8 | 28 | 27 | 8-13 |
| ABIDE - IU | 16 | 4 | 15 | 5 | 17-54 |
| ABIDE - KUL | 28 | 0 | 0 | 0 | 18-35 |
| ABIDE - LEUVEN | 26 | 3 | 30 | 5 | 12-32 |
| ABIDE - OLIN | 17 | 3 | 14 | 2 | 10-24 |
| ABIDE - ONRC | 20 | 4 | 20 | 15 | 18-31 |
| ABIDE - PITT | 26 | 4 | 23 | 4 | 9-35 |
| ABIDE - SBL | 15 | 0 | 15 | 0 | 20-64 |
| ABIDE - STANFORD | 16 | 4 | 16 | 4 | 7-12 |
| ABIDE - TCD | 21 | 0 | 21 | 0 | 10-20 |
| ABIDE - TRINITY | 24 | 0 | 25 | 0 | 12-25 |
| ABIDE - UCD | 14 | 4 | 10 | 4 | 12-17 |
| ABIDE - UCLA | 63 | 7 | 50 | 11 | 7-17 |
| ABIDE - USM | 73 | 2 | 56 | 3 | 8-50 |
| ABIDE - YALE | 20 | 8 | 20 | 8 | 7-17 |
| NIMH | 68 | 17 | 29 | 16 | 1-9 |
| Subtotal | 580 | 80 | 472 | 116 |  |
|  | 660 |  | 588 |  |  |
| Grand Total | 1248 |  |  |  |  |

Demographic information for sites that were excluded from statistical analysis, due either to insufficient females in the autism or TD groups after quality control, or failed segmentation. Site codes are: BNI - Barrow Neurological Institute; CALTECH - California Institute of Technology; CMU - Carnegie Mellon University; EMC - Erasmus University Medical Center Rotterdam; ETH - ETH Zürich; GU - Georgetown University; IU - Indiana University; KUL - Katholieke Universiteit Leuven; LEUVEN - University of Leuven; OLIN - Olin, Institute of Living at Hartford Hospital; ONRC - Olin Neuropsychiatry Research Center, Institute of Living at Hartford Hospital; PITT - University of Pittsburgh School of Medicine; SBL - Social Brain Lab, BCN NIC UMC Groningen and Netherlands Institute for Neurosciences; STANFORD - Stanford University; TCD - Trinity Center for Health Sciences (Release 2); TRINITY - Trinity Center for Health Sciences (Release 1); UCD - University of California, Davis; UCLA - University of California, Los Angeles; USM - University of Utah School of Medicine; YALE - Yale Child Study Center; NIMH - National Institute of Mental Health

### Quality Control and Site Exclusion

Raw scans were inspected visually and rated by two independent raters (SB, ST, and/or MMC), as described by Bedford et al. [4]. Scans with significant motion or other artifacts were processed but were excluded from analysis. Sites with fewer than five females in the autism or TD groups after motion, scan-quality, and segmentation quality control (described below) were excluded from the analysis. This left 1322 individuals across 11 sites in the final dataset. Actual numbers varied according to structure due to variable numbers of segmentation failures, and are detailed in Table 3.

**Table 3:**
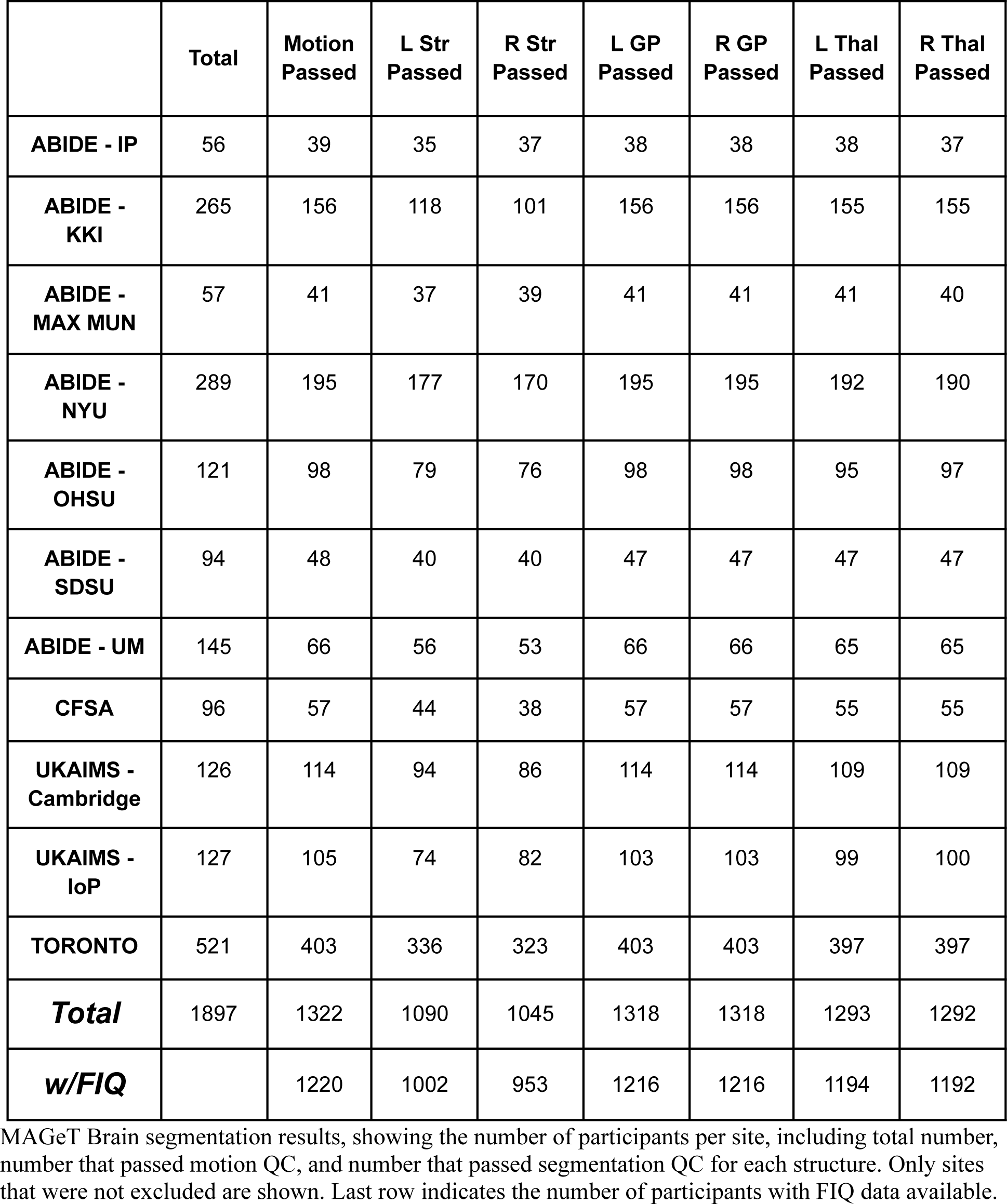
Quality Control Results.

|  | **Total** | **Motion**<br>**Passed** | **L Str**<br>**Passed** | **R Str**<br>**Passed** | **L GP**<br>**Passed** | **R GP**<br>**Passed** | **L Thal**<br>**Passed** | **R Thal**<br>**Passed** |
| --- | --- | --- | --- | --- | --- | --- | --- | --- |
| **ABIDE - IP** | 56 | 39 | 35 | 37 | 38 | 38 | 38 | 37 |
| **ABIDE -**<br>**KKI** | 265 | 156 | 118 | 101 | 156 | 156 | 155 | 155 |
| **ABIDE -**<br>**MAX MUN** | 57 | 41 | 37 | 39 | 41 | 41 | 41 | 40 |
| **ABIDE -**<br>**NYU** | 289 | 195 | 177 | 170 | 195 | 195 | 192 | 190 |
| **ABIDE -**<br>**OHSU** | 121 | 98 | 79 | 76 | 98 | 98 | 95 | 97 |
| **ABIDE -**<br>**SDSU** | 94 | 48 | 40 | 40 | 47 | 47 | 47 | 47 |
| **ABIDE - UM** | 145 | 66 | 56 | 53 | 66 | 66 | 65 | 65 |
| **CFSA** | 96 | 57 | 44 | 38 | 57 | 57 | 55 | 55 |
| **UKAIMS -**<br>**Cambridge** | 126 | 114 | 94 | 86 | 114 | 114 | 109 | 109 |
| **UKAIMS -**<br>**IoP** | 127 | 105 | 74 | 82 | 103 | 103 | 99 | 100 |
| **TORONTO** | 521 | 403 | 336 | 323 | 403 | 403 | 397 | 397 |
| ***Total*** | 1897 | 1322 | 1090 | 1045 | 1318 | 1318 | 1293 | 1292 |
| ***w/FIQ*** |  | 1220 | 1002 | 953 | 1216 | 1216 | 1194 | 1192 |

MAGeT Brain segmentation results, showing the number of participants per site, including total number, number that passed motion QC, and number that passed segmentation QC for each structure. Only sites that were not excluded are shown. Last row indicates the number of participants with FIQ data available.

### Image Processing and Quality Control

All T1-weighted scans, regardless of quality, were preprocessed in site-wise batches, using the minc-bpipe-library pipeline (https://github.com/CoBrALab/minc-bpipe-library). Total brain volume (TBV) was estimated using CIVET 1.1.12 (Montreal Neurological Institute). The thalamus, striatum and globus pallidus were then segmented, their volumes computed, and a surface mesh fit, for each structure in each scan, using the Multiple Automatically Generated Templates (MAGeT Brain) algorithm [25, 26], in batches by site using original subcortical CoBrA Lab atlases as input [27, 28]. All scans at each site were segmented, however only those scans that passed the motion quality control procedure described above were included in the downstream analyses. All labels were inspected visually and rated for segmentation quality [29] by one or more expert raters (DM, and SB, CK, EO, or MMC), and inaccurately labelled structures were excluded from downstream analysis; all successful segmentations were retained. For this reason, the sample size varies between structures, as detailed in Table 3.

### Morphometry

Vertex-wise surface area and displacement, representing convexity or concavity relative to a standard model were computed using MAGeT Brain morphometry across the surfaces of each structure, as described by Tullo et al. [30] and other manuscripts from our group [31–33] and detailed in Supplementary Methods 2.2.

### Statistical Analysis

Given the large number of sites with different scanners, acquisition protocols, and sample demographics, all primary statistical analyses were conducted in a site-wise manner. Within each site, linear regressions were conducted using the lm function in R (4.1.3), from which Hedges *g\** effect sizes and variances were computed [34]. These were then pooled across sites using random-effects meta-analyses [35, 36] using the metafor package (3.4-0) in R (4.1.3), to produce aggregate effect-size measures for the entire dataset. The R code used to generate these meta-analyses is available at https://github.com/dnmacdon/ASD_analysis. The statistical models used are described in Supplementary Methods Section 3.

### Case-Control Analyses

The volumes of the left and right thalamus, globus pallidus, and striatum were modeled using linear regression, and the results were pooled using the meta-analytic technique described above. The models used are given in the Supplementary Methods (6.3.1). In all models, diagnostic status (DX, either autism or TD) was the predictor, and total brain volume (TBV) was included as a covariate. In the initial model, sex and age were also included as covariates. These models were then refit with the addition of FIQ, for the subset of data for which FIQ data was available (Table 3). Statistical analysis of the morphometric data was analogous to that of the volumetric data, except that it was performed at every vertex in each structure, and corrected using false discovery rate (FDR) at 5% [37].

### Volumetric Heterogeneity-Focused Analysis: Sex, Age, and FIQ

To determine whether sex, age, and FIQ are important contributors of volumetric heterogeneity (beyond the effect of total brain volume), a partially nested series of models was fit as described above, to compare models with and without terms for sex, age, and FIQ, and their interactions with diagnosis. The relative fit of these models was estimated using the Akaike Information Criterion (AIC) [38]. Akaike weights were computed from the AIC for each model, which accounts for both goodness of fit and site size, and indicates the strength of evidence in favour of each of the models in a set [39], and weighted according to site size. Because some datasets did not include FIQ information, the analysis was first performed without including FIQ to maximize statistical power, then repeated on the smaller data set with FIQ included. Correction for multiple comparisons was applied after performing the meta-analysis, with an FDR threshold of 5%. For more information, see Supplementary Methods 6.3.3.

### Morphometric Heterogeneity-Focused Analysis: Sex, Age, and FIQ

Vertex-wise models with and without the term of interest were fit by site, the AIC for each was computed, and these were compared pair-wise to determine the best-fitting model at each vertex. These were combined across sites using a winner-take-all approach, weighted by site size, resulting in vertex-wise maps indicating where on the surface of the structure the inclusion of age / sex improved fit. The analysis was then repeated including FIQ. Models are given in Supplementary Methods 6.3.3.

### Follow-up Analyses: Sex-Stratification, and Age/FIQ Centering

Where age or FIQ were found to be important explanatory variables, an age- or FIQ-centered analysis was conducted to evaluate the interaction between autism diagnosis and age/FIQ. This was done by repeating the case-control analysis described above, with an age- or FIQ-by-diagnosis interaction term included in the models and computing the model with age centered (centering value subtracted from all subject ages) at 5-year intervals from ages 6-61, or FIQ centered (centering value subtracted from all subject FIQ) at 10-point intervals from 51-141 (Supplementary Methods 6.3.4). This provided an indication of the effect of autism on structure volume, vertex-wise surface area and displacement at each age interval, without sacrificing statistical power by stratifying the dataset. FDR correction at 5% was done across all vertices, all structures, all age or FIQ intervals, and both measures.

### Confirmatory Analyses: Linear Mixed Effects and ComBat Harmonization

For confirmation of our results and for homology with previous multi-site autism literature, we repeated the case-control analyses using two different techniques that account for intersite differences. First, a mega-analysis was conducted using linear mixed effect models [40], with site included as a random factor. Linear mixed models were computed using the RMINC (v1.5.3.0) package in R [41]. A second mega-analysis was conducted by first harmonizing volume, surface area, and displacement data across sites using ComBat (v1.0.13) in R [42–44], then computing linear regressions on the entire dataset. For details, refer to Supplementary Methods 6.3.2.

### Magnitude of Autism Traits

We assessed the relationship between our subcortical morphometry measures and the level of autistic traits, as measured by the Autism Diagnostic Observation Schedule Calibrated Severity Score (ADOS-CSS). This measure was only available for *n*=239 participants with autism after motion QC (192 male, 47 female), across five sites (KKI, NYU, OHSU, SDSU, TORONTO, UM). The semi-partial correlation between structure-wise volume, vertex-wise surface area, and vertex-wise displacement and ADOS-CSS, while controlling for TBV, age and sex [34] was then computed at each site. These were pooled across sites using random-effects meta-analysis for each structure and vertex. FDR correction at 5% was done across all vertices, all structures, and both measures. For the specific models used, refer to Supplementary Methods 6.3.5.

## Results

### Number of sites and individuals after quality control

Of the 3145 scans over 32 separate sites in the complete dataset, 20 sites, for a total of 1118 participants, were excluded because the sites did not have 5 or more females per group after quality control. The NIMH dataset (130 participants) was also excluded from analysis because of segmentation failure throughout the dataset, perhaps due to the very young ages (range 1-9 years) of the participants, though it was included in previous studies by our group of the cortex [4, 5]. This left 1897 participants across 11 sites. Of these, 1322 (338 M-autism, 496 M-TD, 125 F-autism, 363 F-TD) scans passed motion QC. All of the other excluded sites were in the ABIDE datasets. For details, refer to Tables 1 and 2.

The number of participants varied depending on the specific subcortical structure being examined, as segmentation quality control was performed on a per-structure basis. Common failures included over-segmentation of the caudate nucleus into the third ventricle, and under-segmentation of the anterior caudate nucleus. These failures were reduced by manually correcting atlases, and by using regions of interest to focus nonlinear registration to the specific structure in the specific hemisphere. Quality control results are summarized in Table 3 (improved workflow) and Supplementary Table ST-1 (original workflow).

### No volume differences in autism

We did not observe any significant volume differences in autism vs. TD in the thalamus, striatum, or globus pallidus, when controlling for TBV, age and sex (Figure 1). This was true both when effect sizes were pooled across sites using random-effects meta-analysis, as well as within most sites, with two exceptions: reduced left striatal volume in autism in the ABIDE OHSU site (*p*=.03), and reduced right thalamic volume in autism in the UKAIMS Institute of Psychiatry site (*p=*.04), though even these effects did not survive correction for multiple comparisons (*q*=.66 for both). There were several nearly significant differences at individual sites, including reduced right striatal volume at OHSU (*p*=.07, *q*>.65), increased left striatal volume at UKAIMS Cambridge (*p*=.08, *q*>.65), decreased left striatal volume at ABIDE UM (*p*=.09, *q*>.65), decreased left pallidal volume at IoP (*p*=.08, *q*>.65), increased left and right thalamic volume at ABIDE NYU (*p*=.08, *q*>.65), and decreased left thalamic volume at ABIDE UM (*p*=.07, *q*>.65). The results were similar when the analysis was repeated including FIQ as a covariate: we did not observe any significant volume differences in autism in any of the six structures.

**Figure 1:**
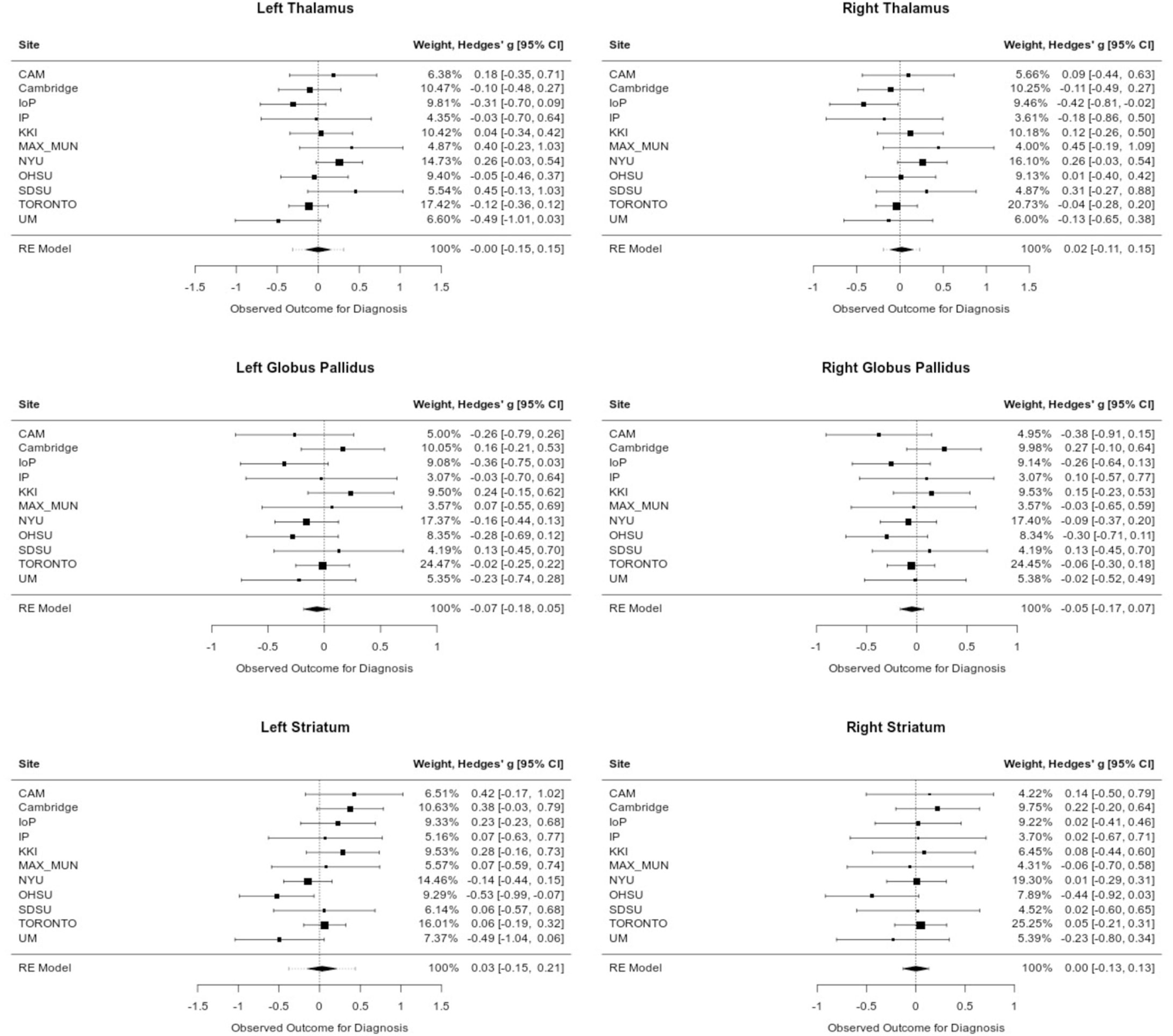
Forest plots showing results of random-effects meta-analysis across sites for all structures. Hedges *g\** effect sizes are reported for main effect of diagnosis in model structure_volume ∼ diagnosis + total_brain_volume + age + sex. No main effect of diagnosis was observed. Columns are: site name, forest plot, site weight, Hedges *g\** estimate and 95% confidence intervals. Site codes are: CAM - Cambridge Family Study of Autism; Cambridge - UKAIMS Cambridge site; IoP - UKAIMS Institute of Psychiatry site; IP - ABIDE Institut Pasteur; KKI - ABIDE Kennedy Krieger Institute; MAX_MUN - ABIDE Ludwig Maximilians University Munich; OHSU - ABIDE Oregon Health and Science University; SDSU - ABIDE San Diego State University; TORONTO - SickKids / University of Toronto; UM - ABIDE University of Michigan

### Role of age, sex, and IQ in volumetric models

Model selection using site size-weighted Akaike weights indicated that age, but not sex, improved model fit for both left and right striatum (Akaike weights 0.46 left, 0.51 right). There was somewhat weaker evidence that both age and sex improved model fit for both left and right thalamus (Akaike weight 0.42 left and right), and weak evidence that model fit was best when neither age nor sex were included for both left and right globus pallidus (Akaike weight 0.35 left, 0.36 right, Supplementary Figure SF-1). We did not observe any effect of autism diagnosis on subcortical volumes in any of the structures when following up with sex-stratified, age-centered, or FIQ-centered analyses.

### Localized effects - alterations of surface area and shape in autism

Localized differences in both surface area and shape (relative displacement) were observed in all structures following vertex-wise analysis (FDR < .05; Figure 2). In the striatum, surface area and displacement effects were limited to the putamen. These mostly comprised a region of areal contraction on the central portion of the right dorsal putamen, and small, mainly anterior regions of areal expansion, as well as a left, anterior region of inward displacement (decreased convexity), near the nucleus accumbens, and two patches of outward displacement on the left antero-ventral and postero-dorsal putamen. We also observed some bilateral areal expansion on the anterior caudate and a very small region of areal contraction on the right dorsal caudate.

**Figure 2:**
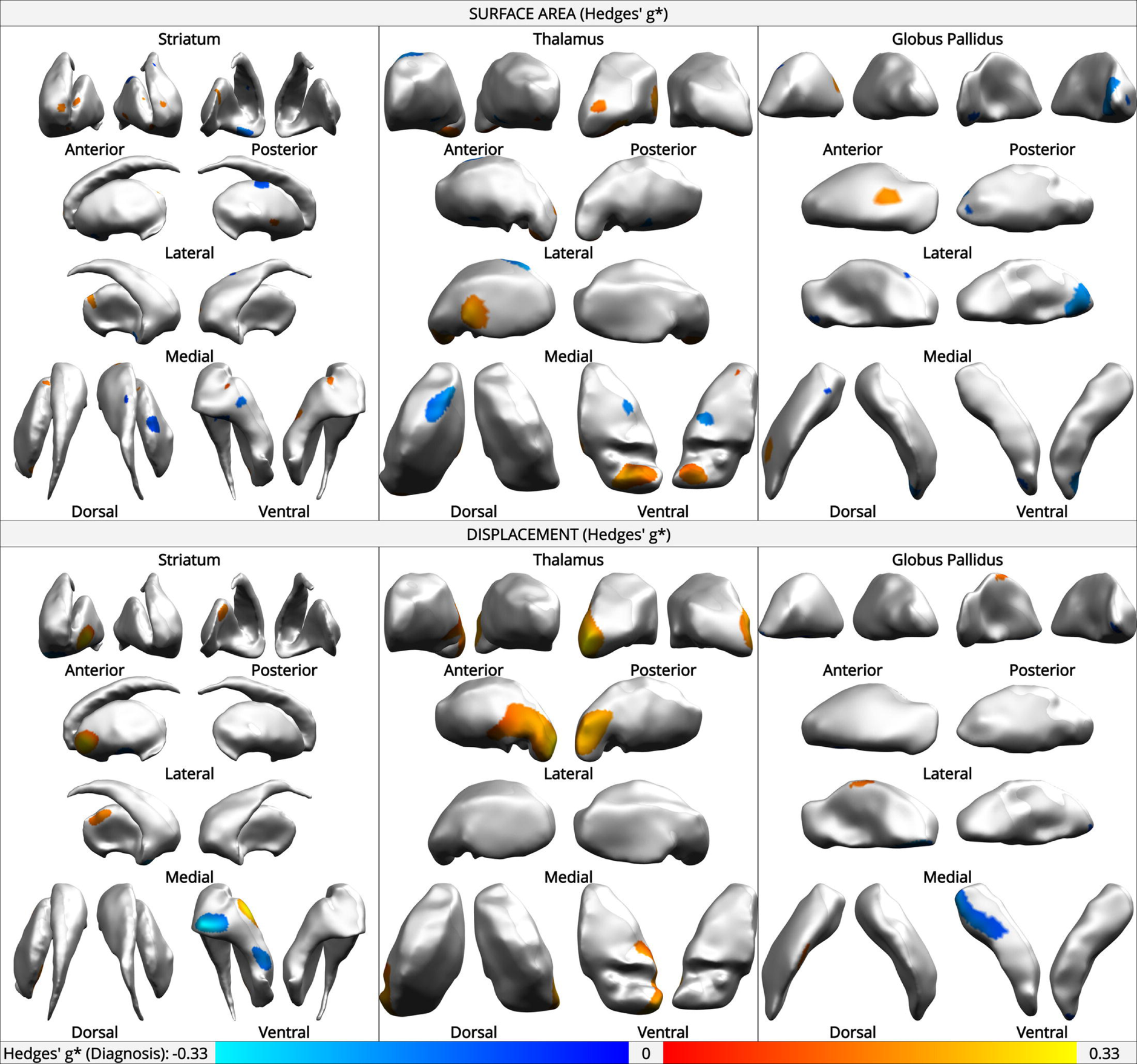
Hedges’ *g\** main effect of autism diagnosis on vertex-wise surface area and displacement in the left and right striatum, thalamus, and globus pallidus, when controlling for total structure volume, age, and sex. Warm colours indicate positive effects, cool colours indicate negative effects (*g\** range −0.3 to 0.3). Surface area is the Voronoi area surrounding a vertex; displacement represents relative convexity (positive) or concavity (negative).

In the thalamus, the largest region of altered surface area was a bilateral region of increase on the ventral posterior surface, approximately corresponding to the ventral surface of the pulvinar. There was also a fairly large region of positive displacement, approximately corresponding with the more lateral surface of the pulvinar and the ventral posterolateral nucleus.

No bilateral effects were observed in the globus pallidus. Surface area effects were mainly a patch of areal expansion on the left lateral surface, a patch of areal contraction on the right posterior medial surface. Displacement effects included a region of positive displacement on the central dorsal medial surface of the left pallidum, and a region of negative displacement on the left anterior ventral surface. Vertices of peak positive and negative effect of diagnosis on surface measures are indicated in Figure 3, with an example breakdown by site of effect sizes and their respective confidence intervals.

**Figure 3:**
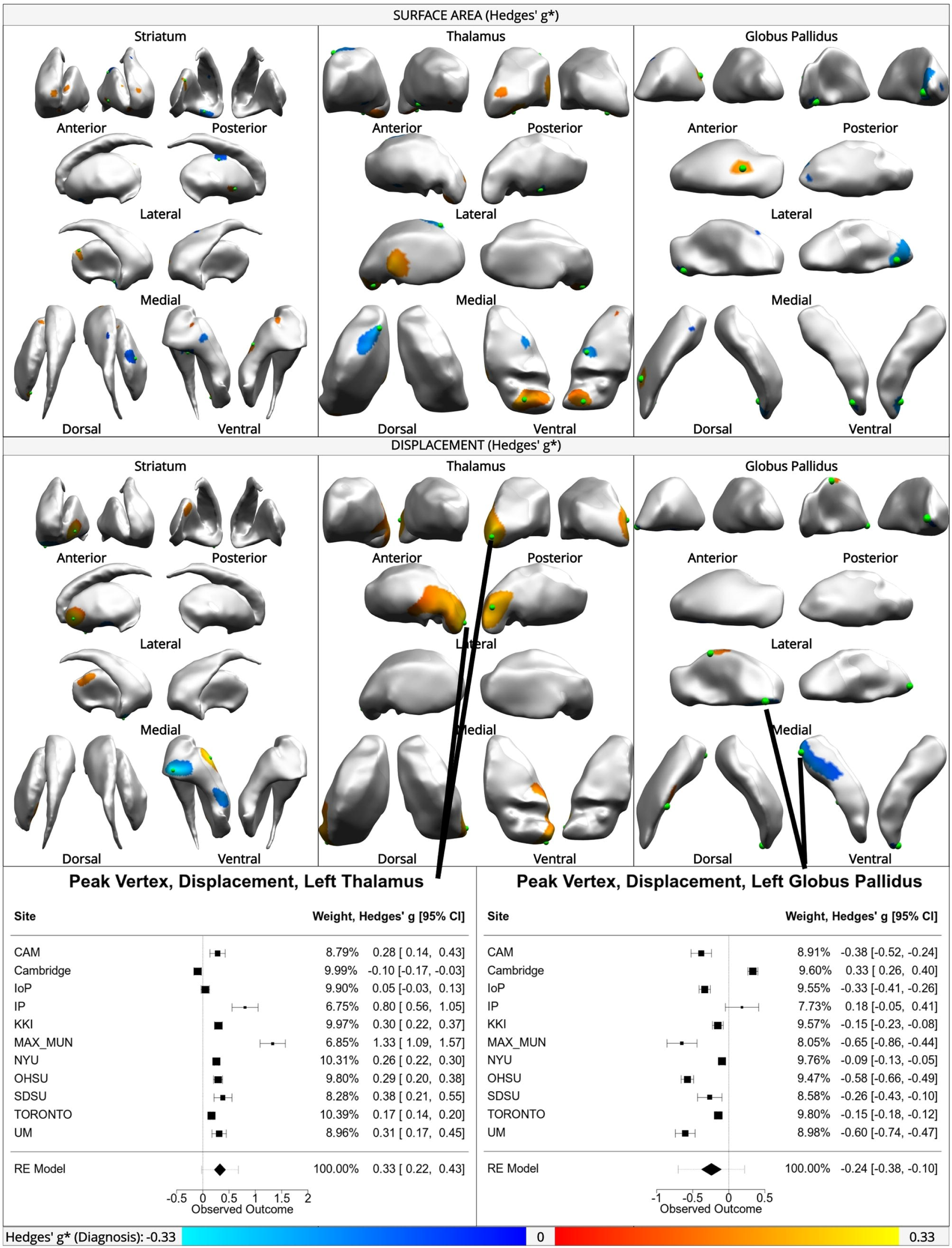
Main effect of diagnosis on vertex-wise surface area (top) and displacement (bottom) in striatum, thalamus, and globus pallidus. Green spheres indicate vertices with peak effect sizes. Forest plots show meta-analytic results at vertices where peak effects were observed for thalamic displacement (positive, left) and pallidal displacement (negative, right).

### Role of age, sex, and FIQ in morphometric models

Vertex-wise model selection analyses indicated that age contributes to the variation in surface-based measures, particularly for displacement, over the surface of most of the globus pallidus, lateral thalamus, and much of the striatum (Supplementary Figure SF-2). Including FIQ does not improve fit when modeling surface area across most structures, with the exception of a small region on the anterior dorsal caudate. There are, however, regions on the surface of all three structures where FIQ improves the fit of models of displacement (Supplementary Figure SF-3). The influence of sex in surface-area models was limited to a relatively small proportion of overall area, with few contiguous regions. Sex was important over somewhat larger regions in all three structures in models of displacement, but these regions still accounted for less than half of overall surface area (Supplementary Figures SF-4, SF-5).

Because of the large proportion of vertices for which age was found to be an important explanatory variable, follow-up age-centered analyses were performed. A representative example is shown in Figure 4, for age-centered thalamic displacement, centered at intervals of five years. This shows large patches of relative increased convexity in autism in the dorso- and medio-lateral right thalamus, as well as a region of the rostro-ventral thalamus roughly corresponding to the pulvinar, but only in childhood. These effects fade by adolescence. In adulthood, relatively little of the thalamus shows any shape effects of autism, though a small region of decreased convexity in autism on the left medial wall appears to be relatively stable throughout adulthood, while there is a patch of relative increased concavity appearing from middle age on, near the left pulvinar region. The remainder of the age-centered results are shown in Supplementary Figures SF-6 through SF-10.

**Figure 4:**
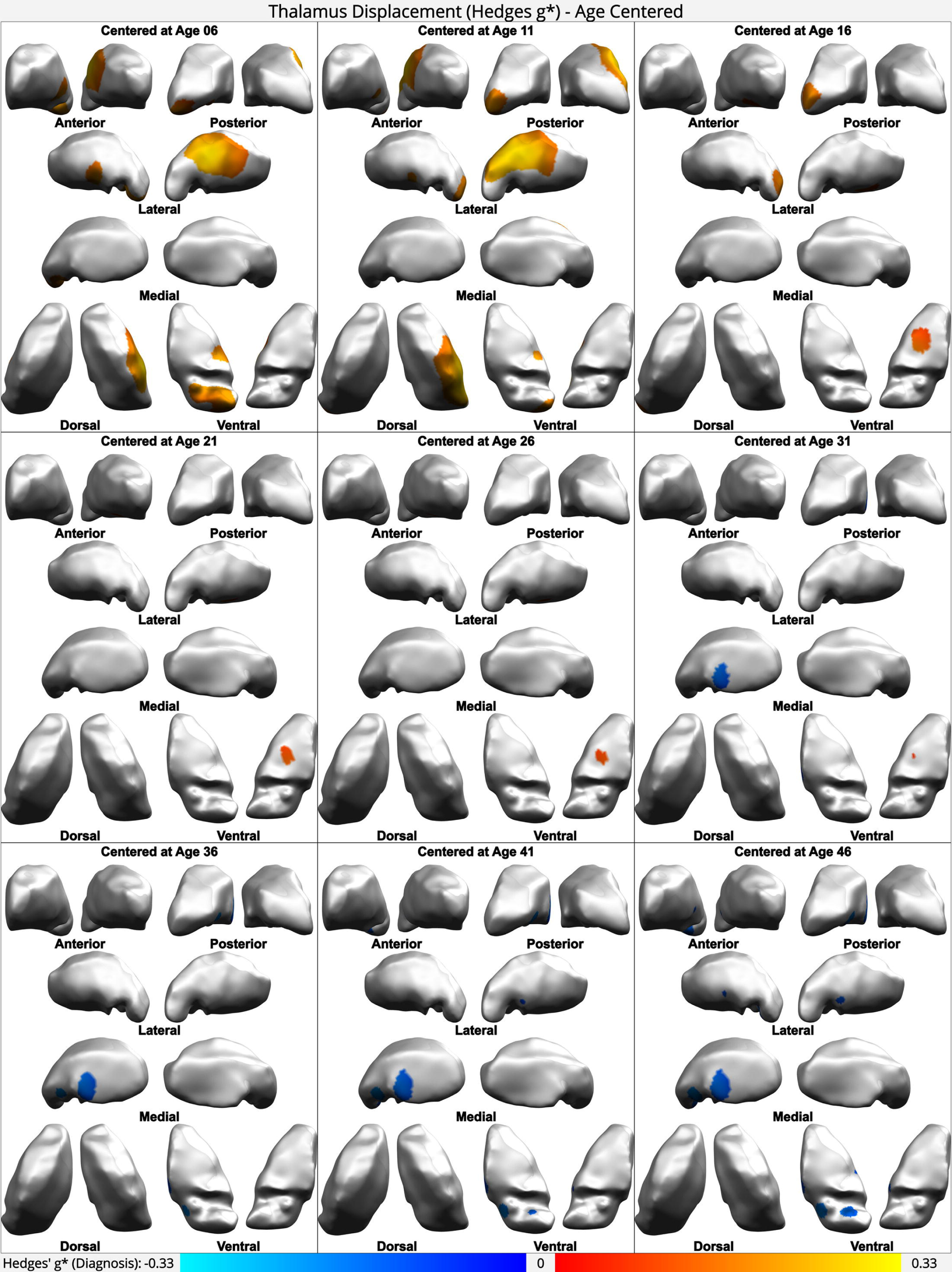
Age-centered analysis, centered on five-year intervals from ages 6-46. Hedges’ *g\** main effect of autism diagnosis on vertex-wise displacement in the left and right thalamus, when controlling for total structure volume and sex. Warm colours indicate positive effects, cool colours indicate negative effects (*g\** range −0.33 to 0.33). Displacement represents relative convexity (positive) or concavity (negative).

### Linear mixed model and ComBat-harmonized mega-analyses confirm results, but are less sensitive

A linear mixed model mega-analysis, including site as a random effect, indicated only two small regions of significant effects of autism diagnosis after multiple comparisons correction, which roughly comprise a subset of the regions of significant effects detected using meta-analysis as described above. These were a region of reduced convexity around the left pulvinar, and a region of areal contraction around the right posterior pole of the globus pallidus. These results are shown in Supplementary Figure SF-11.

Modeling the same data using linear models, after harmonizing across sites using ComBat, while preserving variation due to age and sex, as well as due to age, sex, and FIQ, reproduced the same general patterns, however none of the effects survived correction for multiple comparisons (Supplementary Figures SF-12, SF-13).

### No association between magnitude of autism traits and subcortical volumes or morphometry

No association was found between scores on the ADOS-CSS and subcortical volumes, vertex-wise surface area, or vertex-wise displacement (*p* > .05 for all structures). Only six sites (ABIDE-KKI, ABIDE-NYU, ABIDE-OHSU, ABIDE-SDSU, ABIDE-UM, Toronto) reported ADOS-CSS scores, so this analysis was performed with *n*=241 after motion QC and site exclusion.

## Discussion

In this study we examined volumetric and morphometric differences related to autism in the striatum, globus pallidus, and thalamus in a large, multi-site, cross-sectional MRI dataset containing both males and females, with individuals from 5 to 65 years of age. We found no volumetric differences between the autism and TD groups in any of the structures, but did find several regions of altered surface area and convexity in all three structures. Furthermore, age was an important explanatory variable across more than 50% of all surfaces when considering convexity/concavity, and across more than 50% of the globus pallidus when considering surface area. Sex and FIQ were found to be important explanatory variables across 10-25% of vertices, when considering displacement, and across fewer than 10% of vertices when considering surface area. No association was found between the magnitude of autism traits and volumetric or morphometric measures. Overall, our findings help confirm that the neurodevelopment of the striatum, globus pallidus and thalamus are affected in autism, in subtle ways that are not consistent across space or time.

Our findings underscore the importance of accounting for age when examining neuroanatomical variation associated with autism. While we did not observe volumetric differences in any of the structures across the lifespan, we did find a complex pattern of spatially localized morphometric differences that were highly dependent upon age. Normative studies have shown that, in typically developing individuals, the thalamus, striatum, and globus pallidus do not undergo uniform growth, but rather show complex patterns of localized growth and contraction through childhood, adolescence, and adulthood [32, 45]. It may be that localized expansion and contraction in different regions has a cancellation effect, such that detectable volume changes may be subtle. These growth patterns also differ between structures, and change over the lifetime. The volume of the striatum, for instance, tends to peak during childhood or adolescence and then decrease monotonically throughout adulthood, whereas thalamic volume can remain stable for decades [45–47], though the timing of peak volume attainment can differ between the sexes [32].

Our findings of localized surface area and displacement group differences were more pronounced in childhood and adolescence while attenuated and in different regions in adulthood. In the context of normative development, the regional effects we found did not coincide with areas of significant areal expansion or contraction with age in normative development through adulthood. In a study of normative subcortical shape and volume changes across the adult lifespan from ages 18-83 years, Tullo et al. [45] reported general volumetric decline in all three structures, as well as localized regions of areal contraction in the anterior medial and ventral posterior thalamus, the bilateral medial wall of the globus pallidus, and the nucleus accumbens and the medial wall of the putamen, as well as areal expansion with age in the tail of the caudate nucleus. None of these regions coincided with the regions of differential areal contraction or expansion in autism that we observed over the lifespan. The picture is somewhat less clear during childhood and adolescence, where large swathes of each structure show contraction or expansion with age, particularly in the thalamus and caudate body and tail [32].

Surprisingly, there was a lack of concordance between our morphometric results and those of a study that used similar techniques and a dataset that also included ABIDE. Schuetze et al. (2016) reported localized morphometric differences in autism in the thalamus, globus pallidus and striatum that were not reflected in structure volumes. However, the regions in which they observed an effect of autism on surface area or shape did not in general coincide with the regions that we report here. Aside from some differences in methodology, including quality control and data combination techniques, their sample included only male participants 35 years of age and younger. Some regions where they reported differences overlap with regions where we found that sex made a significant contribution to model accuracy (Supplementary Figure SF-4). This suggests that sex may be an important modulator within those regions.

While the under-representation of females in autism studies is improving, autism is continually understudied in females, despite evidence that sex modulates neuroanatomy in both autism [4, 6] and normative development of subcortical structures [32]. Consequently, there is very little evidence regarding sex differences in autism in the structures studied here. One recent large study performed by the ENIGMA consortium found no sex-by-diagnosis interaction effect on the volumes of any of these structures [22], which is consistent with our results. The picture is less clear when looking at localized morphometry. Including sex in the model improves the fit in 5-10% of vertices when evaluating surface area, and 15-25% of vertices for relative displacement (Supplemental Figure SF-5). In particular, the medial caudate bilaterally, and much of the dorsomedial thalamus, when considering the effect of autism on localized displacement, may qualitatively exhibit sex differences (Supplemental Figure SF-4), despite the lack of evidence for a sex-by-diagnosis interaction in their volumes. This corresponds to the findings of a recent study using a very different methodology on a dataset that overlaps with ours by including data from the Toronto SickKids site [48]. Normative development of these structures through childhood and adolescence diverges somewhat between the sexes: Raznahan et al. [32] reported regions of sexually differential areal expansion and contraction in the striatum, globus pallidus, and thalamus. Nevertheless, there is very little concordance between these regions of normative sexually differential maturation and the regions we found were differentially affected by sex in autism.

FIQ is also often unexamined in studies of autism, despite evidence of FIQ-associated variation in subcortical neuroanatomical differences. Various studies have found correlations between FIQ and regional volumes of both cortical and subcortical structures [49–51]. We did not evaluate this directly in the structures under study, but rather asked whether accounting for FIQ improves our modelling of subcortical anatomy in autism. We did not find any evidence to this effect, which is consistent with several reports [9, 22].

We used the ADOS-CSS as a measure of the magnitude of autism traits, to evaluate the relationship with our measures of subcortical volumes and morphometry. Unfortunately, there was no consistent measure available in all of the datasets: a variety of ADOS versions and modules were used, resulting in *n*=241 participants. Our finding of no relationship in any of the structures is at odds with the findings of other groups, which have found associations between restricted, repetitive behaviours and pallidal surface area [9], as well as growth rates of the caudate [52]. However, our divergence from those studies may be due to the relatively low statistical power in this portion of the study.

A number of limitations of this study should be considered. First, the data was compiled from many smaller datasets, each collected for different purposes. There is no harmonization between datasets in terms of sample characteristics, inclusion/exclusion criteria, MRI hardware, software, or acquisition parameters, or behavioural measures captured. To allow for meaningful statistical analysis, we used a meta-analytic technique that has been used successfully before [4, 5], and followed up with two other commonly used data combination techniques: linear mixed effect models and ComBat harmonization. Our meta-analytic technique proved to be more sensitive than both of these other common strategies, as we have shown previously at the cortical level [4]. The ENIGMA OCD working group performed a similar comparison at the cortical level and reached the opposite conclusion, though their study had several methodological differences, including the fact that cortical statistics were examined by Desikan-Killiany region [53] rather than vertex-wise, and quality control was automated [54]. As discussed above, a region-wise analysis may not be sufficiently sensitive to subtle effects that do not neatly overlap with a predetermined regional map. Also, it has been shown that quality control decisions can significantly influence the order and even direction of effects [55]. For this reason, we visually inspected each scan and each segmentation, and adopted a strict quality-control protocol.

A related issue is the paucity of behavioural measures that were available across multiple sites, and the lack of other potentially relevant demographic information, such as socioeconomic status. There is some evidence that accounting for such behavioural and demographic heterogeneity improves the sensitivity of surface-based morphological measures, at least in small samples [11]. Large datasets that include consistent behavioural measures, such as the Quebec 1000 Families cohort now underway (q1k.ca), will make possible large studies that account for behavioural heterogeneity in a more comprehensive way.

The cross-sectional nature of the data also limits to some degree the scope of interpretation of these results. While some datasets did include longitudinal scans, for the majority of participants only a single time point was available. This makes it difficult to draw any direct conclusions about how neurodevelopmental trajectories may be altered in autism. Also, although the dataset includes participants from ages 5 to 65 years, it is heavily weighted towards the younger end of the age range.

In addition, after quality control and removal of sites with too few females to allow for statistical analysis, we were only able to retain between 31% and 40% of participants, depending on the structure. This is an unfortunately high rate of data attrition, however, given that including relatively poor quality scans can introduce artifactual effects [4, 55], it was a necessary step. That said, while the effect of motion on cortical measures is well documented, particularly at distal regions such as the orbitofrontal cortex and temporal poles, the effect on the subcortical morphometric measures used here are currently less well understood and, considering their central location in the brain, may not be as drastic.

Finally, this study relies on a case-control analysis, comparing at the group level the neuroanatomy of those who have been diagnosed with autism with those who have not. The underlying assumption is that there are meaningful, group-wise differences in neuroanatomy. While this may be the case, the overwhelming discord in the literature discussed in the Introduction, and the highly space- and time-variable nature of our findings, may call this assumption into question. Recent studies using techniques that can account for individual variation, such as subtyping and normative modelling [56], have begun to appear in the autism literature, and provide another lens through which to examine autism-related neuroanatomical differences.

Overall, our findings help confirm that the neurodevelopment of the striatum, globus pallidus and thalamus are altered in autism, in a subtle location-dependent manner that is not reflected in overall structure volumes, and that is highly non-uniform across the lifespan.

## Acknowledgements

This research was funded in part by the National Science and Engineering Research Council and the Fonds de Recherche du Québec - Santé, in the form of graduate student funding to DNM. SAB received a graduate student fellowship from the Healthy Brains, Healthy Lives initiative of McGill University, funded by the Canada First Research Excellence Fund. The Autism Imaging Multicentre Study Consortium was funded by the Medical Research Council United Kingdom grant G0400061. The Cambridge Family Study of Autism was funded by a Clinical Scientist Fellowship from the UK Medical Research Council (MRC) (G0701919). AR was supported by funding from the Intramural Research Program of the NIMH (Clinical trial reg. no. NCT00001246, clinicaltrials.gov; NIH Annual Report Number, ZIA MH002794, Protocol ID 89-M-0006). The Toronto sample was gathered from studies supported by grants MOP-119541, MOP-106582 and MOP-14237 from the Canadian Institutes of Health Research (to MT), and from the POND Network, funded by the Ontario Brain Institute (grant IDS-I l-02 to EA and JL), an independent non-profit corporation, funded partially by the Ontario government. The opinions, results and conclusions are those of the authors and no endorsement by the Ontario Brain Institute is intended or should be inferred. JL received funding from the Canadian Institute for Health Research. MMC received funding from the Canadian Institute for Health Research, the Natural Sciences and Engineering Research Council, the Fonds de recherche du Québec – Santé and McGill University’s Healthy Brains for Healthy Lives initative. SBC was supported by the Autism Research Trust. DGM was supported in this work by funding from the MRC UK, the National Institute for Health Research (NIHR) Biomedical Research Centre at South London and Maudsley NHS Foundation Trust, and King’s College London (Medical Research Council grant no. G0400061 to DGMM). DGM, SBC, AR, JS, CE, and RH are also supported by EU-AIMS and AIMS-2 TRIALS. EU-AIMS receives support from the Innovative Medicines Initiative (IMI) Joint Undertaking (JU) under grant agreement no. 115300, the resources of which are composed of financial contributions from the European Union’s Seventh Framework Programme (grant FP7/2007-2013). AIMS-2 TRIALS received support from EFPIA and AUTISM SPEAKS, Autistica, and SFARI, and funding from the IMI 2 JU under grant agreement no. 777394, with support from the European Union’s Horizon 2020 research and innovation programme. MVL was supported by an ERC Starting Grant (ERC-2017-STG; 755816). M-CL was supported by the O’Brien Scholars Program within the Child and Youth Mental Health Collaborative at the Centre for Addiction and Mental Health and the Hospital for Sick Children, Toronto, the Slifka-Ritvo Award for Innovation in Autism Research by the Alan B. Slifka Foundation and the International Society for Autism Research, and the Academic Scholars Award from the Department of Psychiatry, University of Toronto, and the Canadian Institutes of Health Research Sex and Gender Science Chair (GSB 171373). The authors would like to thank the investigators and participants in the ABIDE dataset. Funding sources for each individual sites are provided on the official ABIDE website (http://fcon_1000.projects.nitrc.org/indi/abide/).

