## Supplemental Materials for "Characterizing Subcortical Structural Heterogeneity in Autism"

**Supplementary Material**

### 1. Datasets[1]

**ABIDE I & ABIDE II.** The Autism Brain Imaging Data Exchange (ABIDE) is a large-scale, multisite publicly available database consisting of structural and functional magnetic resonance imaging (MRI) as well as behavioural and phenotypic data from individuals with autism and controls.[2, 3] The ABIDE initiative includes two releases of data, ABIDE I and II, including structural MRI data from 1101 and 1044 subjects, respectively (aged 5-64 years), amounting to a total of 1017 individuals with ASD (137 female/880 male) and 1128 controls (274 female/854 male). These data are collected from more than 24 sites internationally (with a number of sites contributing data to both ABIDE I and II). Sites with no females were excluded from our analysis, as were sites which did not have enough females left after quality control (see Quality Control section for details). For information about specific sites and scanning acquisition protocols, see the ABIDE website : http://fcon_1000.projects.nitrc.org/indi/abide/.

**NIMH**. The sample contributed by the National Institute of Mental Health (NIMH) is a longitudinal dataset consisting of 192 T1-weighted MRIs from 85 children with ASD (17 Female/68 Male) and 45 controls (16 Female/29 Male), aged 2-9 years. Each individual had 1 or 2 structural MRI scans. Due to the young age of participants within this dataset, there were more failures due to motion. Therefore, to maximise data, all scans were processed and for individuals with multiple images, if both images passed motion QC, the scan with the better CIVET output was used. If these were also of equivalent quality, the earlier image was used. Diagnosis of ASD was confirmed using the Autism Diagnostic Observation Schedule (ADOS) and the Autism Diagnostic Interview-Revised (ADI-R). Children with ASD were scanned under sedation to minimise movement during scanning, while controls were scanned during natural sleep, or while watching a movie. Scanning was conducted on a 1.5 T General Electric Signa scanner (Milwaukee, WI), using a 3D spoiled gradient recalled echo sequence, with contiguous 1.5 mm axial slices. Image acquisition parameters were as follows: Echo time, 5 ms; repetition time, 24 ms; flip angle 45°; acquisition matrix, 256 3 192; number of excitations, 1; and field of view, 24 cm, resulting in a voxel resolution of 0.9375×0.9375.0×1.5 mm (for details, see Smith et al[4] and Raznahan et al[5]). This study was approved by an NIH Institutional Review Board, and written informed consent was obtained from all participants' parents/guardians.

**UK MRC-AIMS**. The UK Medical Research Council (MRC) Autism Imaging Multicentre Study (AIMS) consists of structural MRI data from 159 adults with ASD (62 F/97 M), and 154 controls (59 F/95 M), aged 18-52 years, collected at the Institute of Psychiatry, Psychology and Neuroscience (IoPPN), King's College London (n = 136); the Autism Research Centre, University of Cambridge (n = 138); and the Autism Research Group, University of Oxford (n = 39). As the University of Oxford sample contained only males, only data from the IoPPN and Cambridge were included in this study. ASD diagnosis was confirmed using the ADI-R. Scanning was conducted using a 3T GE Medical Systems HDx scanners with an 8-channel receive-only RT head-coil. A Driven Equilibrium Single Pulse Observation of T1 (DESPOT1)

[1] The datasets (Section 1) and pre-processing steps (Section 2.1) used in this study were previously assembled and performed by our group, and were first described by Bedford et al. [1]. The dataset descriptions from that report are reproduced here verbatim.

sequence was used to standardise structural MRI scans across sites[6–8], in which quantitative T1 maps are used to create simulated T1-weighted inversion recovery images, with 1×1×1 mm resolution, field of view of 25 cm, repetition time of 1800 ms, inversion time of 50 ms, and flip angle of 20° (for details, see Lai et al[9], Ecker et al[10], Zeestraten et al[11]). This study was approved by the National Research Ethics Committee, Suffolk, England, and written informed consent was obtained from all participants. Despite this standardization of acquisition sequences, in our analyses, Cambridge and the IoPPN were treated as separate sites to minimise any possibilities of site-specific effects.

**Cambridge Family Study of Autism**. The Cambridge Family Study of Autism (CFSA) consists of data from 56 individuals with ASD (17 F/39 M), 40 siblings of individuals with ASD (28 F/12 M), and 40 controls (20 F/20 M), aged 12-18 years. For the purposes of this study, only data from individuals with ASD and controls were analysed. ASD diagnosis was confirmed using the Autism Diagnostic Observation Schedule-Generic (ADOS-G) and the ADI-R. All individuals were recruited by the Autism Research Centre, University of Cambridge. Magnetization prepared rapid gradient echo (MPRAGE) scans were acquired using a 3T Siemens Tim Trio scanner (Siemens Healthcare, Germany) at the MRC Cognition and Brain Sciences Unit (CBU) in Cambridge, with 1 × 1 × 1 mm resolution, repetition time of 2250 ms, echo time of 2.98 ms, inversion time of 900 ms and flip angle of 9° (for more detail see Holt et al[12] and Ypma et al[13]). This study was approved by the Cambridgeshire 1 Research Ethics Committee, and all subjects and/or their parent(s) provided written consent.

**Hospital for Sick Children**. A longitudinal sample of 131 individuals with ASD (25 Female/106 Male) and 390 controls (194 Female/196 Male), aged 4-65 years, were recruited and scanned at the Hospital for Sick Children, Toronto, Ontario, Canada. Like the NIMH sample, the data acquisition for this sample was longitudinal: individuals had between 1-3 structural MRI images. However, we chose to use slightly different QC inclusion criteria since the data were from older participants and therefore more images were expected to pass QC for motion. For participants with multiple scans, the best scan after quality control for motion was selected. If scans were of equivalent quality, the earlier scan was used. ASD diagnosis was confirmed using the ADOS-G and/or ADI-R. Imaging was conducted on a 3T Siemens Trio MRI scanner (MAGNETOM Tim Trio, Siemens AG, Erlangen, Germany), with a 12-channel head coil. A T1-weighted MPRAGE sequence was used, with 1 × 1 × 1 mm resolution, repetition time of 2300 ms, echo time of 2.96 ms, inversion time of 900 ms and flip angle of 9°. Participants were scanned while watching a movie, and foam padding was used to stabilise their heads to restrict motion. This study was approved by The Hospital for Sick Children Research Ethics Board, and written informed consent was obtained from all participants and/or their parent(s).

#### 2 Image Processing

##### 2.1 Preprocessing and Total Brain Volume Estimation

Raw scans were inspected visually and rated by two independent raters, as described by Bedford et al. (2019). Scans with significant motion or other artefacts were processed but were excluded from analysis. All scans, regardless of quality, were preprocessed. Preprocessing was performed in site-wise batches, using the minc-bpipe-library pipeline [14]. This pipeline applies bias field correction, to remove artefactual, low-frequency inhomogeneity in image intensities, using the iterative N4 algorithm [15]. Next, brain masks were generated and used to extract the brains from surrounding tissue (skull, fat, etc.), using the Brain Extraction based on nonlocal Segmentation Technique [16]. Total brain volume was estimated using CIVET 1.1.12 (Montreal Neurological Institute), by summing estimates of grey matter volume, white matter volume, and cerebrospinal fluid volume.

Adjustments were made to the preprocessing pipeline for some sites to maximize the amounts of usable data; the site-based, meta-analytic nature of the statistical analysis ensures that we did not introduce new site-based confounds by doing this. Scans from ABIDE I and II releases from New York University (NYU) were processed with blood vessel masking activated in CIVET. The UK-AIMS scans were subjected to an intensity standardization relative to the MNI ICBM 152 average brain template [17] using minc_nyul [18]. Scans from both ABIDE releases from the Kennedy Krieger Institute (KKI) were pre-processed without brain masking or extraction. Scans from the ABIDE II release from the Institut Pasteur (IP) without bias field correction.

##### 2.2 Segmentation with MAGeTbrain

Segmentation of the thalamus, striatum and globus pallidus was performed using the Multiple Automatically Generated Templates (MAGeT Brain) algorithm [19, 20], in batches by site. MAGeT Brain makes use of five atlases in which voxels of the structures of interest are labeled. These labels were derived from serial histological data, fit to a single high-resolution MRI template [21], then warped to five new high-resolution MRI atlas templates [22]. The 5 label sets were then propagated to a set of 21 template scans, drawn from the data in each batch, giving 105 label sets for each structure. These labels were then propagated to each scan in the batch, producing 105 candidate labels for each structure and for each scan in the dataset. These were merged into a final set of labels for each structure and each scan through a voxel-wise majority vote. Label propagation from one image to another was performed at each step using the Advanced Normalization Tools (ANTs) library [23], by first performing an affine registration, then a non-linear registration between the two volumes, and then using the generated transforms to warp the labels into the space of the second volume. The volume of each structure, for each participant, was computed using the number of voxels multiplied by acquisition specific voxel volume in the label for that structure and the voxel volume.

While other software exists to label and compute the volume of subcortical structures, including FreeSurfer (https://surfer.nmr.mgh.harvard.edu), and FSL-FIRST (https://fsl.fmrib.ox.ac.uk), MAGeT Brain has been shown to compute labels for these structures that more closely resemble the gold standard of manually generated labels created by expert raters [24]. All scans at each site were segmented, however only those scans that passed the motion quality control procedure described above were included in the downstream analyses.

To improve segmentation quality after the initial run, two modifications were made to this workflow. First, to reduce oversegmentation of the striatum, which was a common failure, atlases with manually corrected striatal segmentations were used. Second, segmentation was performed on left and right hemispheres separately, and a subcortical mask was applied to remove the cortex from the registration process. Only the results of this improved image-processing workflow are presented.

#### 3 Statistical Analysis

##### 3.1 Case-control comparisons

The basic models below were used for all case-control analyses, both volumetric and vertex-wise. "struct" represents the left or right striatum, thalamus, or globus pallidus, and "i" represents each vertex across a given structure.

$$V_{struct} = \beta_0 + \beta_1 Diagnosis + \beta_2 TBV + \beta_3 Age + \beta_4 Sex + \varepsilon_{struct}$$

$$SA_i = \beta_{0(i)} + \beta_{1(i)} Diagnosis + \beta_{2(i)} TBV + \beta_{3(i)} Age + \beta_{4(i)} Sex + \varepsilon_i$$

$$Disp_i = \beta_{0(i)} + \beta_{1(i)} Diagnosis + \beta_{2(i)} TBV + \beta_{3(i)} Age + \beta_{4(i)} Sex + \varepsilon_i$$

These models were refit with the addition of FIQ for the subset of data for which FIQ data was available:

$$V_{struct} = \beta_0 + \beta_1 Diagnosis + \beta_2 TBV + \beta_3 Age + \beta_4 Sex + \beta_5 FIQ + \varepsilon_{struct}$$

$$SA_i = \beta_{0(i)} + \beta_{1(i)} Diagnosis + \beta_{2(i)} TBV + \beta_{3(i)} Age + \beta_{4(i)} Sex + \beta_{5(i)} FIQ + \varepsilon_i$$

$$Disp_i = \beta_{0(i)} + \beta_{1(i)} Diagnosis + \beta_{2(i)} TBV + \beta_{3(i)} Age + \beta_{4(i)} Sex + \beta_{5(i)} FIQ + \varepsilon_i$$

All of these models were fit within each site, and Hedges' *g** (an unbiased form of Cohen's *d,* [25]) was computed from the $\beta_1$ coefficient, capturing the effect size of the main effect of diagnosis. These site-wise main effects were then pooled across sites using random effects meta-analysis using the metafor package in R [26].

##### 3.2 Confirmatory analyses

Volumetric and vertex-wise results obtained using the basic case-control models were confirmed using two forms of mega-analysis: linear mixed models, and ComBat harmonization followed by multiple linear regression. Linear mixed models were computed using the RMINC (1.5.3.0) package in R [27], and were specified as follows, using a random intercept term for site:

$$V_{struct} = \beta_0 + \beta_1 Diagnosis + \beta_2 TBV + \beta_3 Age + \beta_4 Sex + \beta_5 FIQ + (1|site) + \varepsilon_{struct}$$

$$SA_i = \beta_{0(i)} + \beta_{1(i)} Diagnosis + \beta_{2(i)} TBV + \beta_{3(i)} Age + \beta_{4(i)} Sex + \beta_{5(i)} FIQ + (1|site) + \varepsilon_i$$

$$Disp_i = \beta_{0(i)} + \beta_{1(i)} Diagnosis + \beta_{2(i)} TBV + \beta_{3(i)} Age + \beta_{4(i)} Sex + \beta_{5(i)} FIQ + (1|site) + \varepsilon_i$$

For the second confirmatory analysis, ComBat harmonization was performed with the neuroCombat (1.0.13) library for R [28–30], prior to multiple linear regression using the basic case-control comparison models described above. Structure volumes were harmonized with respect to site, using the default options, including using parametric priors as well as empirical Bayes-based shrinkage of location and scale parameters across features. For volumetric data, the six structures (left and right globus pallidus, striatum and thalamus) comprised the features. For vertex-wise data, the surface area or displacement at each vertex comprised the set of features. TBV, Sex, and Age were included as biological covariates, to protect against the removal of variance explained by these variables.

##### 3.3 Sex, Age, and FIQ: Heterogeneity-Focused Analysis

For the volumetric analysis, a series of nested multiple linear regression models was fit within each site, with the "global" model including as covariates: total brain volume, age and its interaction with diagnosis, and sex with its interaction with diagnosis. Subsequent models deleted age and its interaction, sex and its interaction, or both. AIC was computed for each model, and these were used to compute Akaike weights for the set of models at each site [31]. These were weighted by site size and averaged across sites to provide a single summary value indicating the relative degree of evidence in favour of each model. This analysis was conducted without FIQ to maximize statistical power, then repeated with FIQ included with the individuals for whom FIQ data was provided in the dataset. For volumetric models, the role of sex, age, and FIQ was evaluated using the following set of models:

*Global:*
$V_{struct} = \beta_0 + \beta_1 Diagnosis + \beta_2 TBV + \beta_3 Age + \beta_4 Sex + \beta_5 Age*Diagnosis + \beta_6 Sex*Diagnosis + \varepsilon_{struct}$

*Age:*
$V_{struct} = \beta_0 + \beta_1 Diagnosis + \beta_2 TBV + \beta_3 Age + \beta_5 Age*Diagnosis + \varepsilon_{struct}$

*Sex:*
$V_{struct} = \beta_0 + \beta_1 Diagnosis + \beta_2 TBV + \beta_4 Sex + \beta_6 Sex*Diagnosis + \varepsilon_{struct}$

*TBV:*
$V_{struct} = \beta_0 + \beta_1 Diagnosis + \beta_2 TBV + \varepsilon_{struct}$

*DX:*
$V_{struct} = \beta_0 + \beta_1 Diagnosis + \varepsilon_{struct}$

For the vertex-wise analyses, the same series of nested models was fit and AIC was computed, at each vertex and within each site. The AIC of the model containing all terms was compared against that of the model with the term of interest deleted (age or sex), and the model with the lower AIC was deemed to be a better fit. These were combined across sites in a winner-take-all approach, weighted by site size, and then mapped over the surface of each structure, giving one map for age and one for sex, indicating locations on the surface where the variable of interest improved model fit. The proportion of vertices for which age and sex improved model fit was computed for each structure. This analysis was then repeated with FIQ included. For vertex-wise models, comparisons of AIC were made between the following pairs of models at each vertex: Global vs. Age, Global vs. Sex, Global vs. FIQ.

##### 3.4 Age- and FIQ-Centered Analysis

The models used in this analysis were as follows for age, where $X_j$ indicates the intervals on which ages were centered. Analogous models were used for FIQ-centering.

$$V_{struct} = \beta_0 + \beta_1 Diagnosis + \beta_2 TBV + \beta_3(Age\text{-}X_j) + \beta_4(Age\text{-}X_j)*Diagnosis + \beta_5 Sex + \varepsilon_{struct}$$

$$SA_i = \beta_{0(i)} + \beta_{1(i)}Diagnosis + \beta_{2(i)}TBV + \beta_{3(i)}(Age\text{-}X_j) + \beta_{4(i)}(Age - X_j) + \beta_{5(i)}Sex + \varepsilon_i$$

$$Disp_i = \beta_{0(i)} + \beta_{1(i)}Diagnosis + \beta_{2(i)}TBV + \beta_{3(i)}(Age\text{-}X_j) + \beta_{4(i)}(Age - X_j) + \beta_{5(i)}Sex + \varepsilon_i$$

##### 3.5 Magnitude of Autism Traits

The ADOS-2 Calibrated Severity Score (CSS) was used as a measure of the magnitude of autism traits. This measure was available for *n*=239 participants with ASD after motion QC (192 male, 47 female), across five sites (KKI, NYU, OHSU, SDSU, TORONTO, UM).

Site-wise models used were as follows:

$$V_{struct} = \beta_0 + \beta_1 CSS + \beta_2 TBV + \beta_3 Age + \beta_4 Sex + \varepsilon_{struct}$$

$$SA_i = \beta_{0(i)} + \beta_{1(i)}CSS + \beta_{2(i)}TBV + \beta_{3(i)}Age + \beta_{4(i)}Sex + \varepsilon_i$$

$$Disp_i = \beta_{0(i)} + \beta_{1(i)}CSS + \beta_{2(i)}TBV + \beta_{3(i)}Age + \beta_{4(i)}Sex + \varepsilon_i$$

The semi-partial correlation [25] was then computed from $\beta_1$, and these were pooled across sites using random-effects meta-analysis as in the base case-control analysis, for each structure and vertex.

#### Supplementary Figures

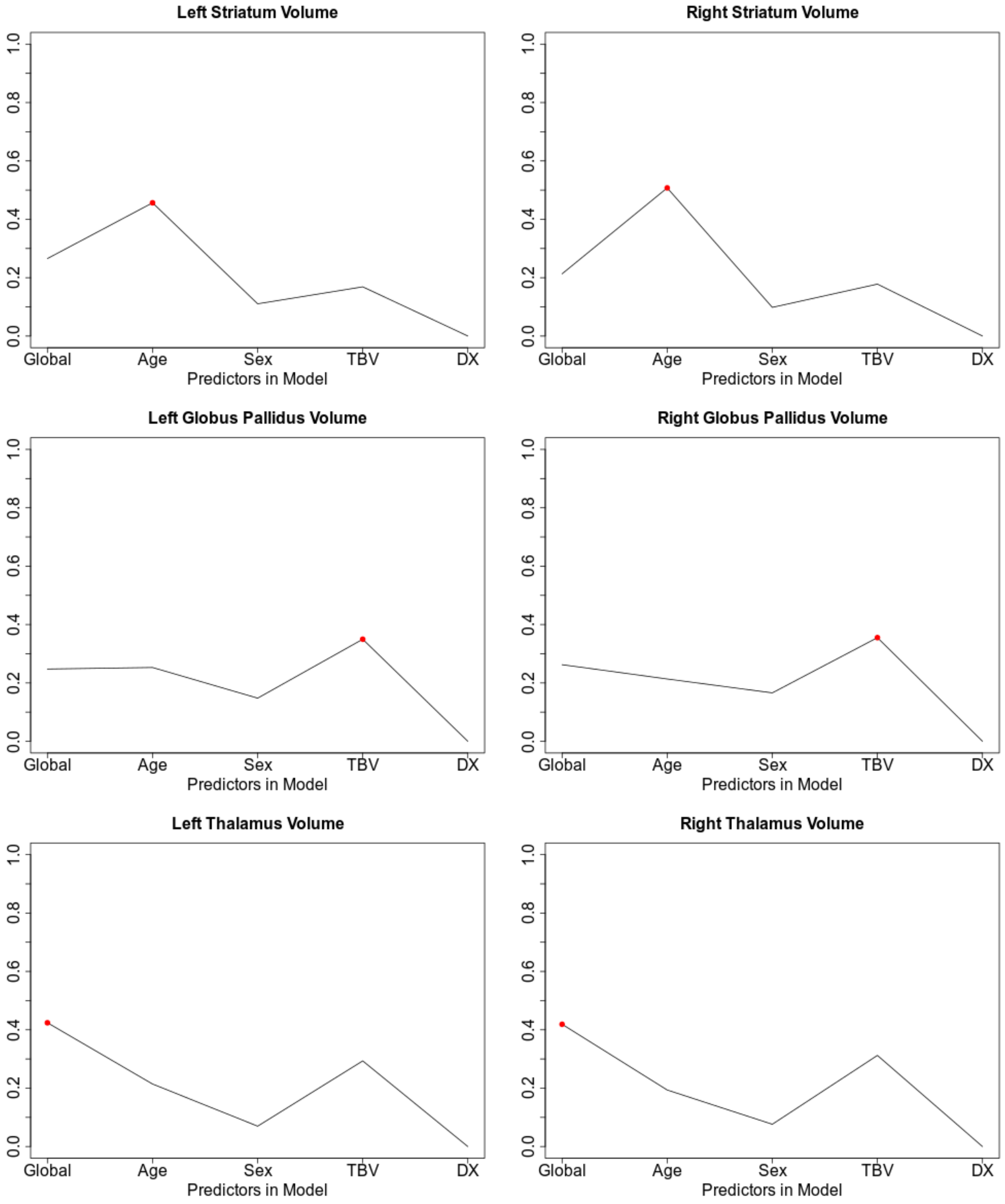


**Supplementary Figure SF-1:** Site-size weighted average of Akaike weights showing evidence for each of five candidate models for left and right striatum, globus pallidus, and thalamus. Red dots indicate maximum value across candidate models. Models are: DX (volume ~ diagnosis), TBV (volume ~ diagnosis + TBV), Sex (volume ~ diagnosis + TBV + Sex + DX*Sex), Age (volume ~ diagnosis + TBV + Age + diagnosis*Age), Global (volume ~ diagnosis + TBV + Age + diagnosis*Age + Sex + diagnosis*Sex)

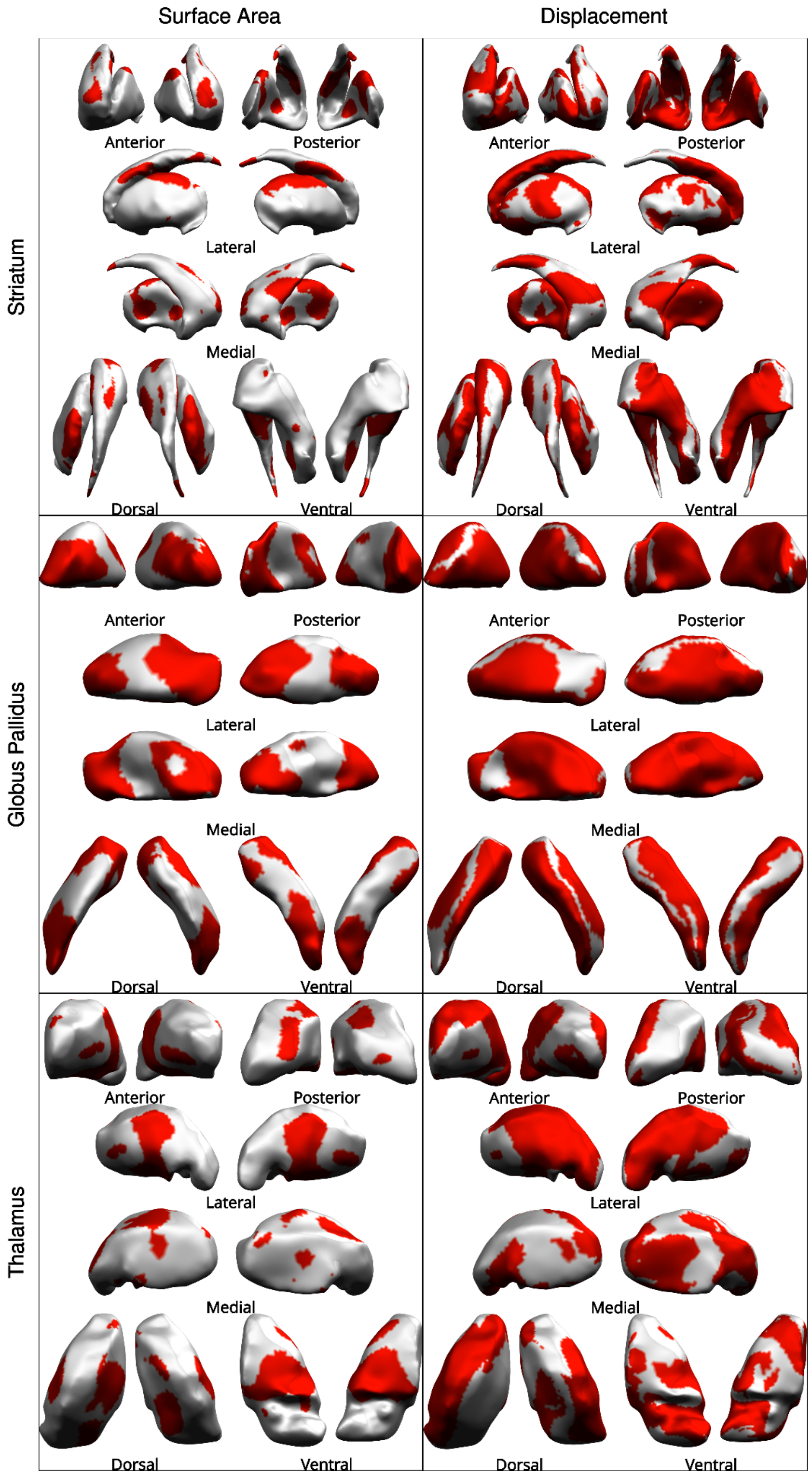


**Supplementary Figure SF-2:** Age maps: Regions on the surface of left and right striatum, thalamus, and globus pallidus in which the inclusion of age and age-by-diagnosis terms improved model fit, for linear models of surface area and displacement on diagnosis, when controlling for sex. Red indicates improved fit with age terms.

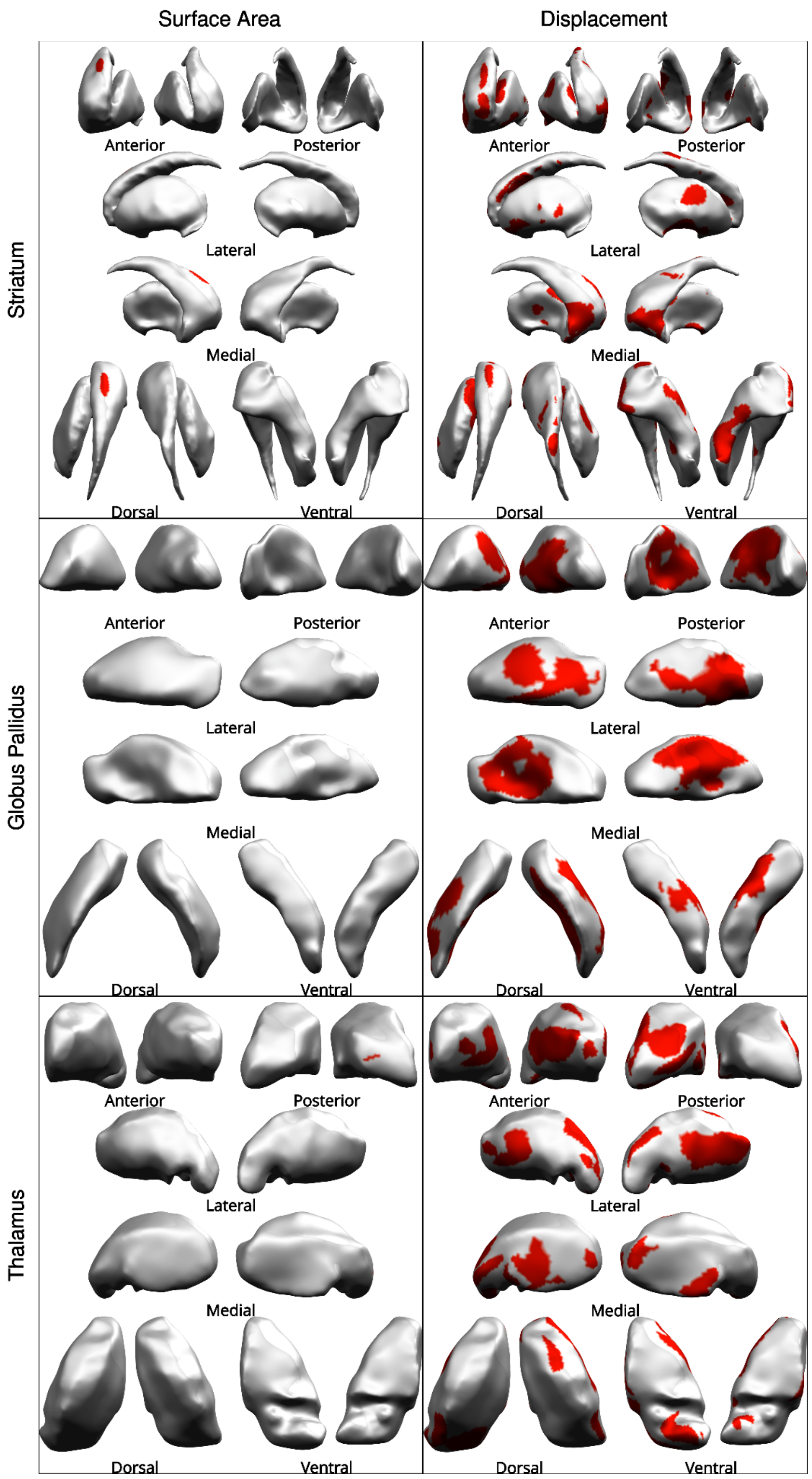


**Supplementary Figure SF-3:** FIQ maps: Regions on the surface of left and right striatum, thalamus, and globus pallidus in which the inclusion of FIQ and FIQ-by-diagnosis terms improved model fit, for linear models of surface area and displacement on diagnosis, controlling for age and sex. Red indicates improved fit with FIQ.

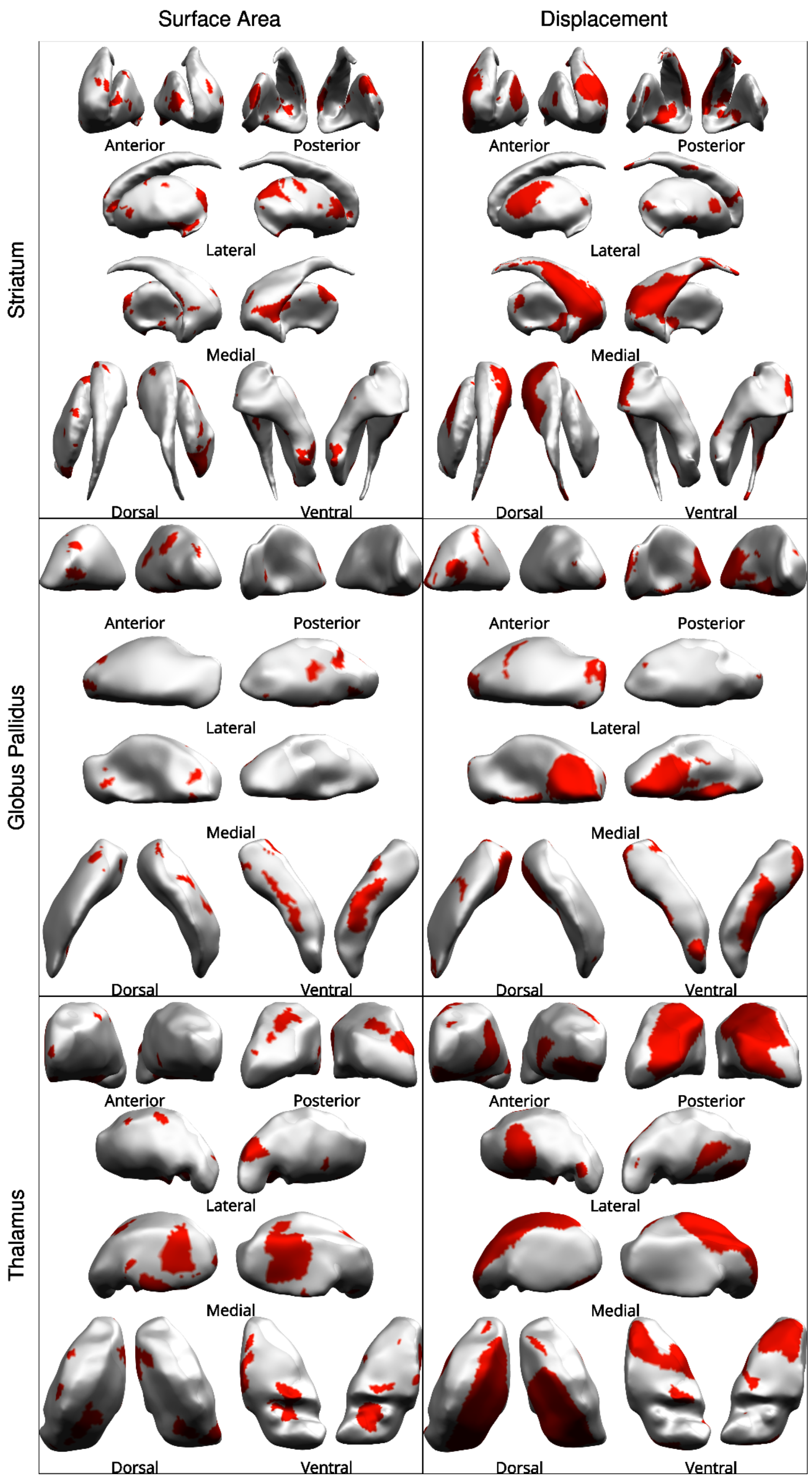


**Supplementary Figure SF-4:** Sex maps: Regions on the surface of left and right striatum, thalamus, and globus pallidus in which the inclusion of sex and sex-by-diagnosis terms improved model fit, for linear models of surface area and displacement on diagnosis, when controlling for age. Red indicates improved fit with sex terms.

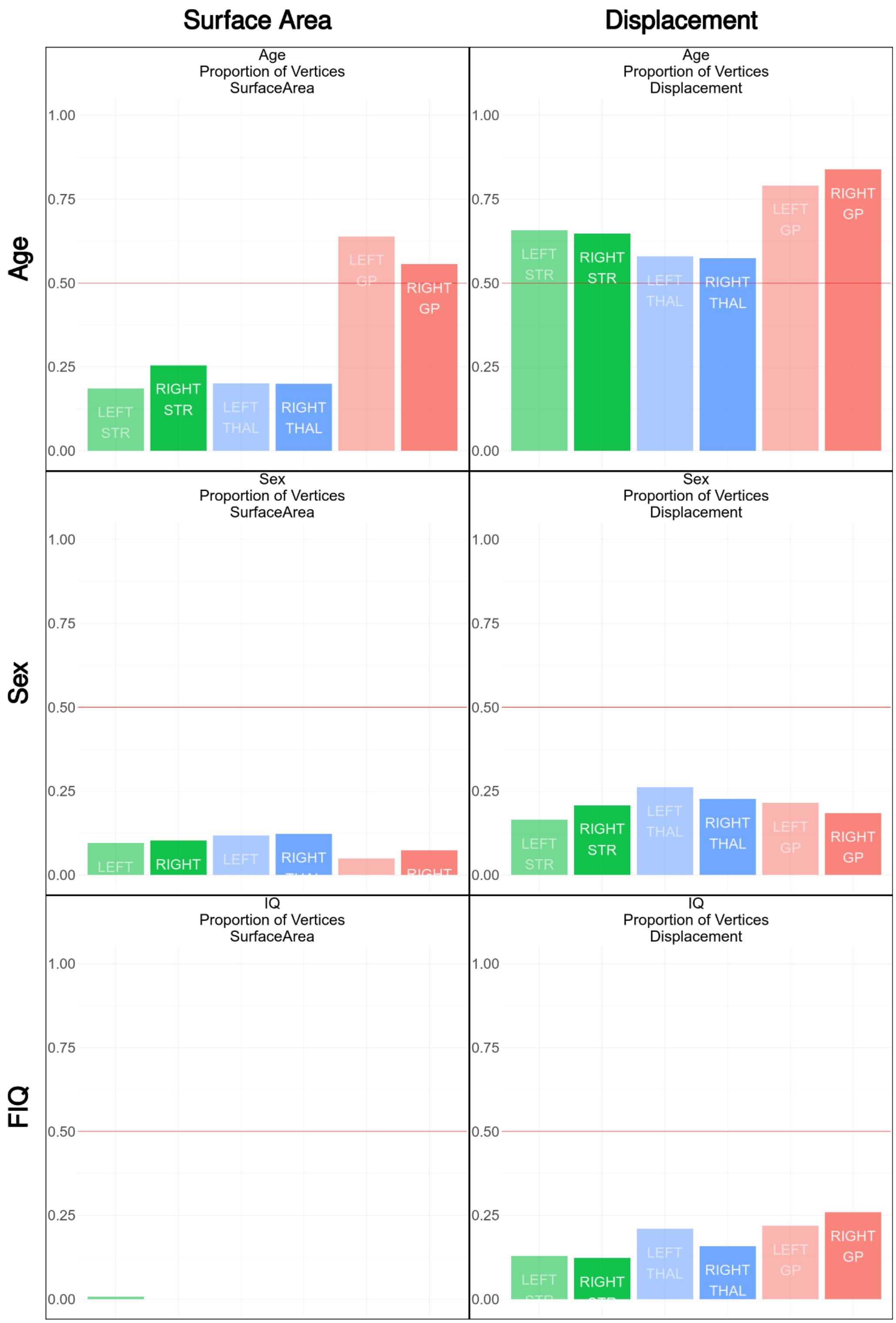


**Supplemental Figure SF-5**: Proportion of vertices for which sex, age, and FIQ improved model fit across left (light) and right (dark) striatum (green), thalamus (blue), and globus pallidus (red). Linear models of main effect of ASD diagnosis on vertex-wise surface area (left column) and displacement (right column). Red line indicates 50%.

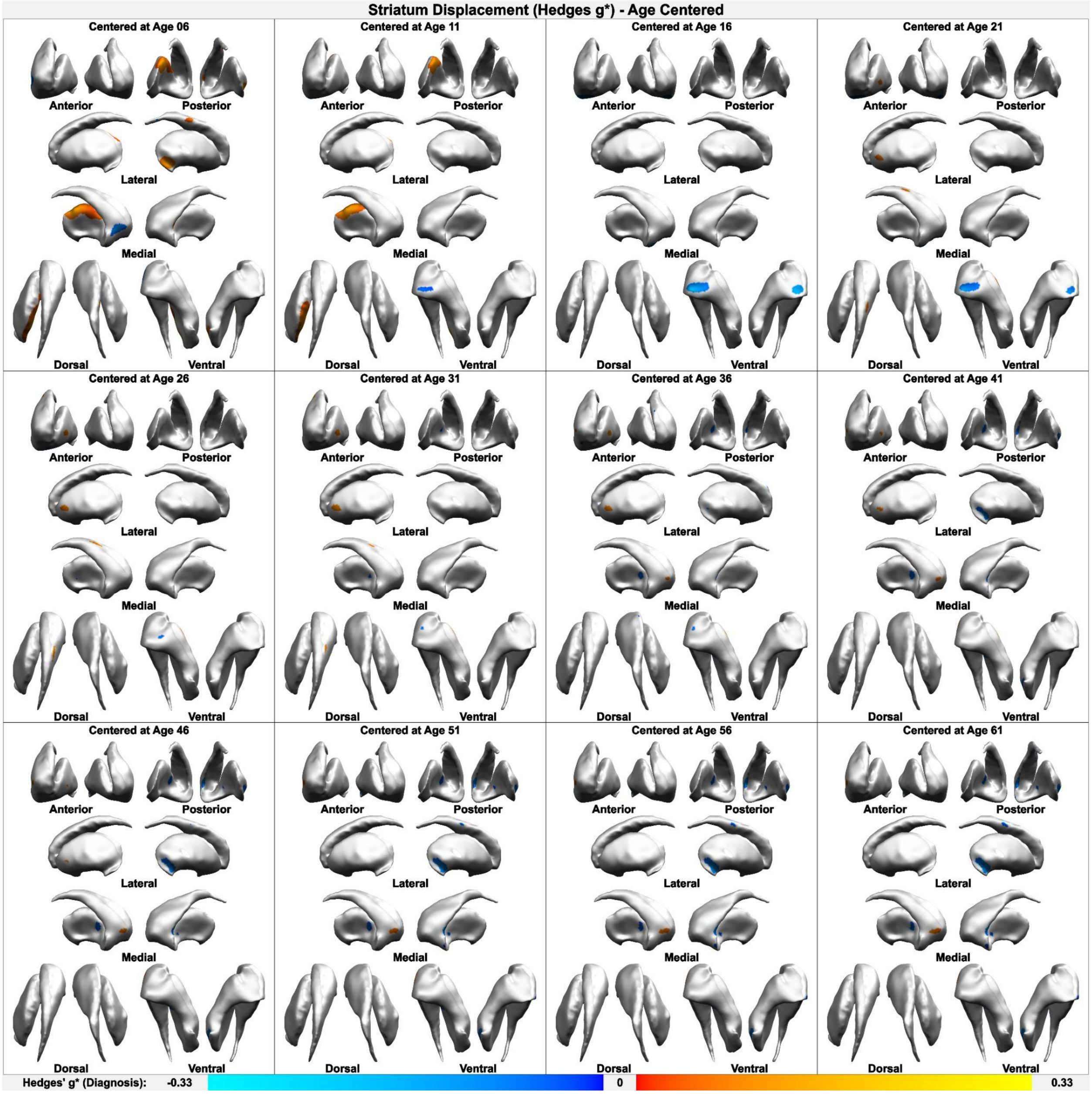


**Supplementary Figure SF-6**: Age-centered analysis, centered on five-year intervals from ages 6-61. Hedges' $g*$ main effect of ASD diagnosis on vertex-wise displacement in the left and right striatum, when controlling for total structure volume and sex. Warm colours indicate positive effects, cool colours indicate negative effects ($g*$ range -0.33 to 0.33). Displacement represents relative convexity (positive) or concavity (negative).

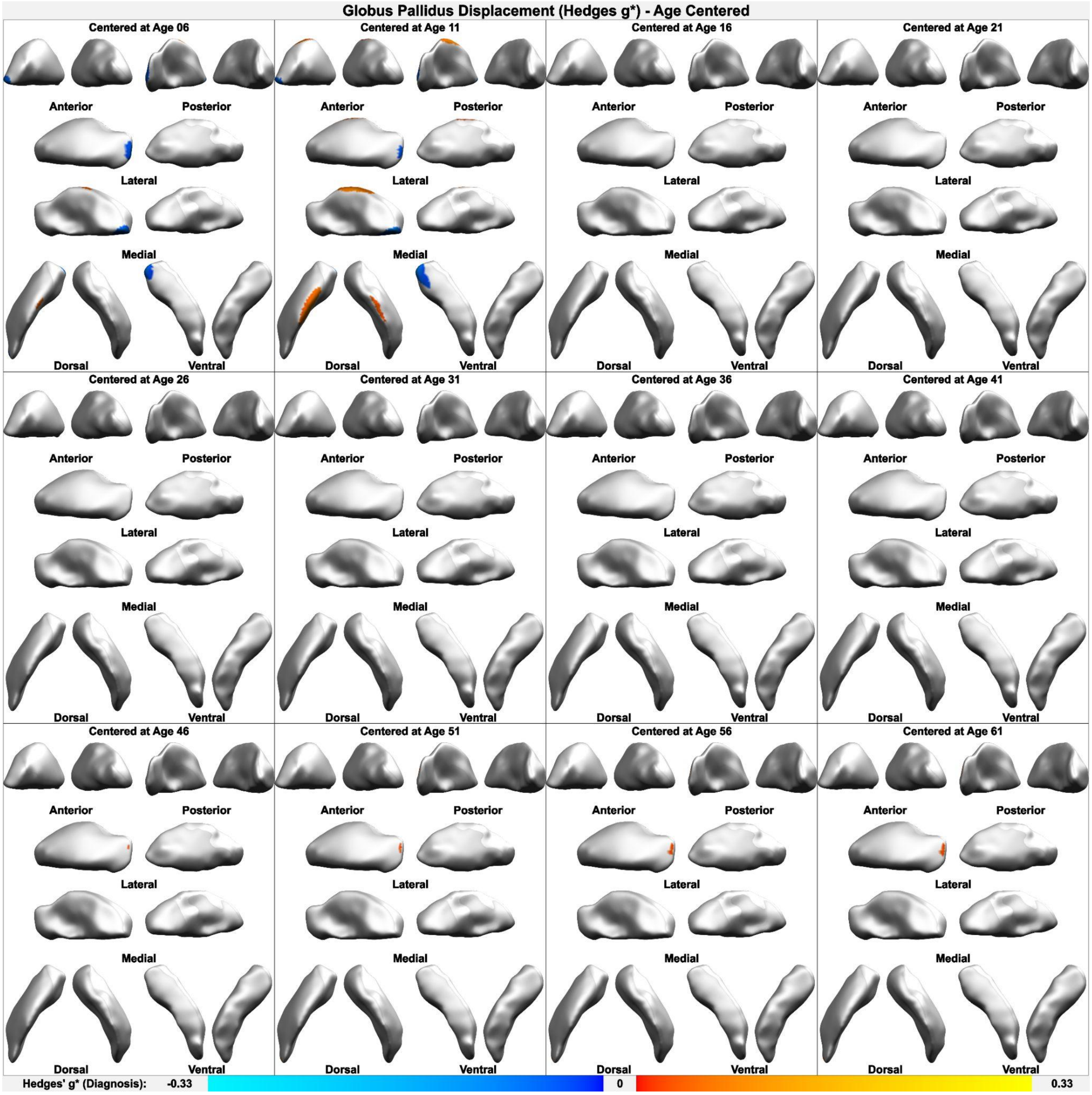


**Supplementary Figure SF-7**: Age-centered analysis, centered on five-year intervals from ages 6-61. Hedges' $g*$ main effect of ASD diagnosis on vertex-wise displacement in the left and right globus pallidus, when controlling for total structure volume and sex. Warm colours indicate positive effects, cool colours indicate negative effects ($g*$ range -0.33 to 0.33). Displacement represents relative convexity (positive) or concavity (negative).

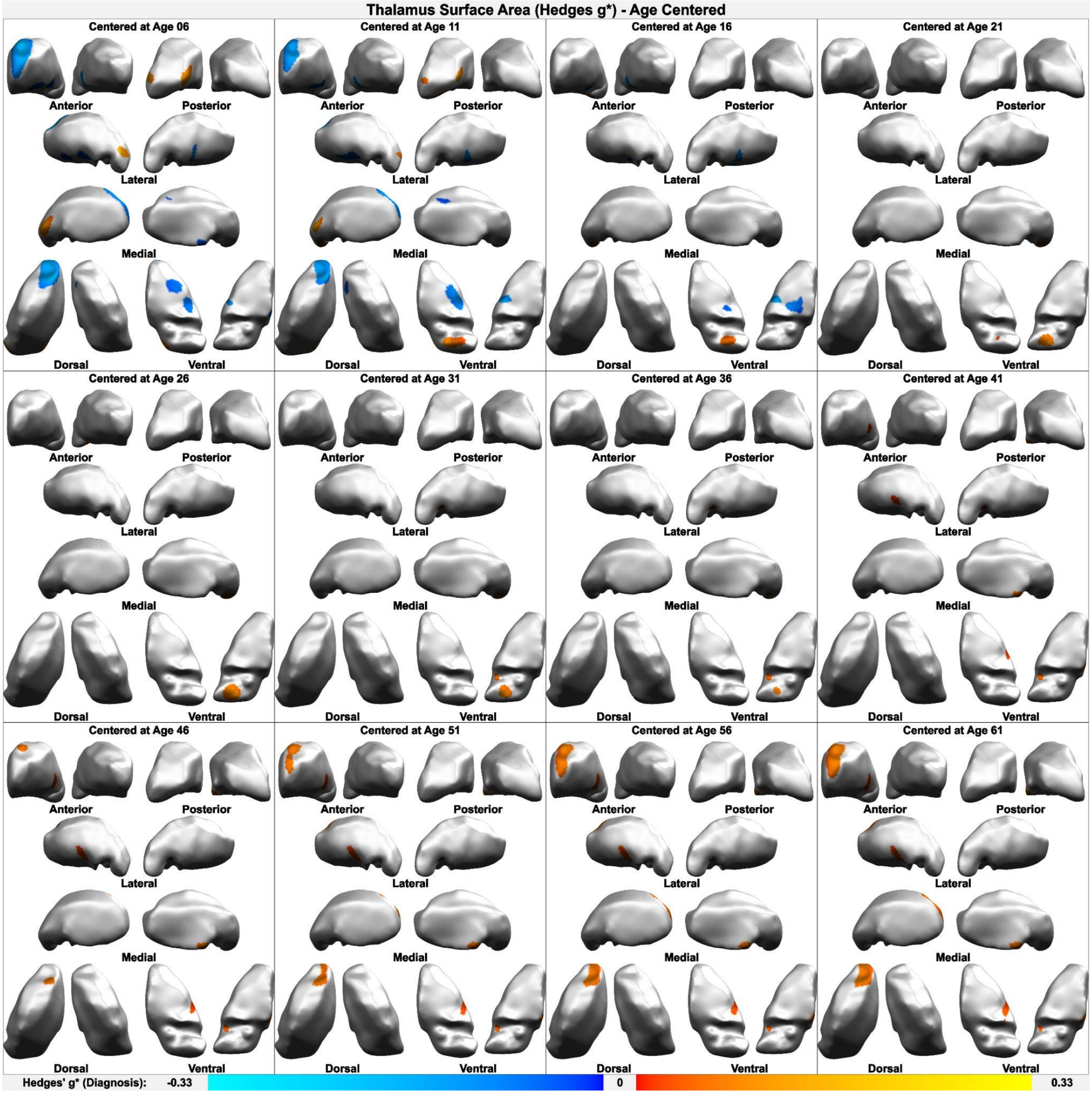


**Supplementary Figure SF-8**: Age-centered analysis, centered on five-year intervals from ages 6-61. Hedges' $g^*$ main effect of ASD diagnosis on vertex-wise surface area in the left and right thalamus, when controlling for total structure volume and sex. Warm colours indicate positive effects, cool colours indicate negative effects ($g^*$ range -0.33 to 0.33). Surface area is the Voronoi area surrounding a vertex.

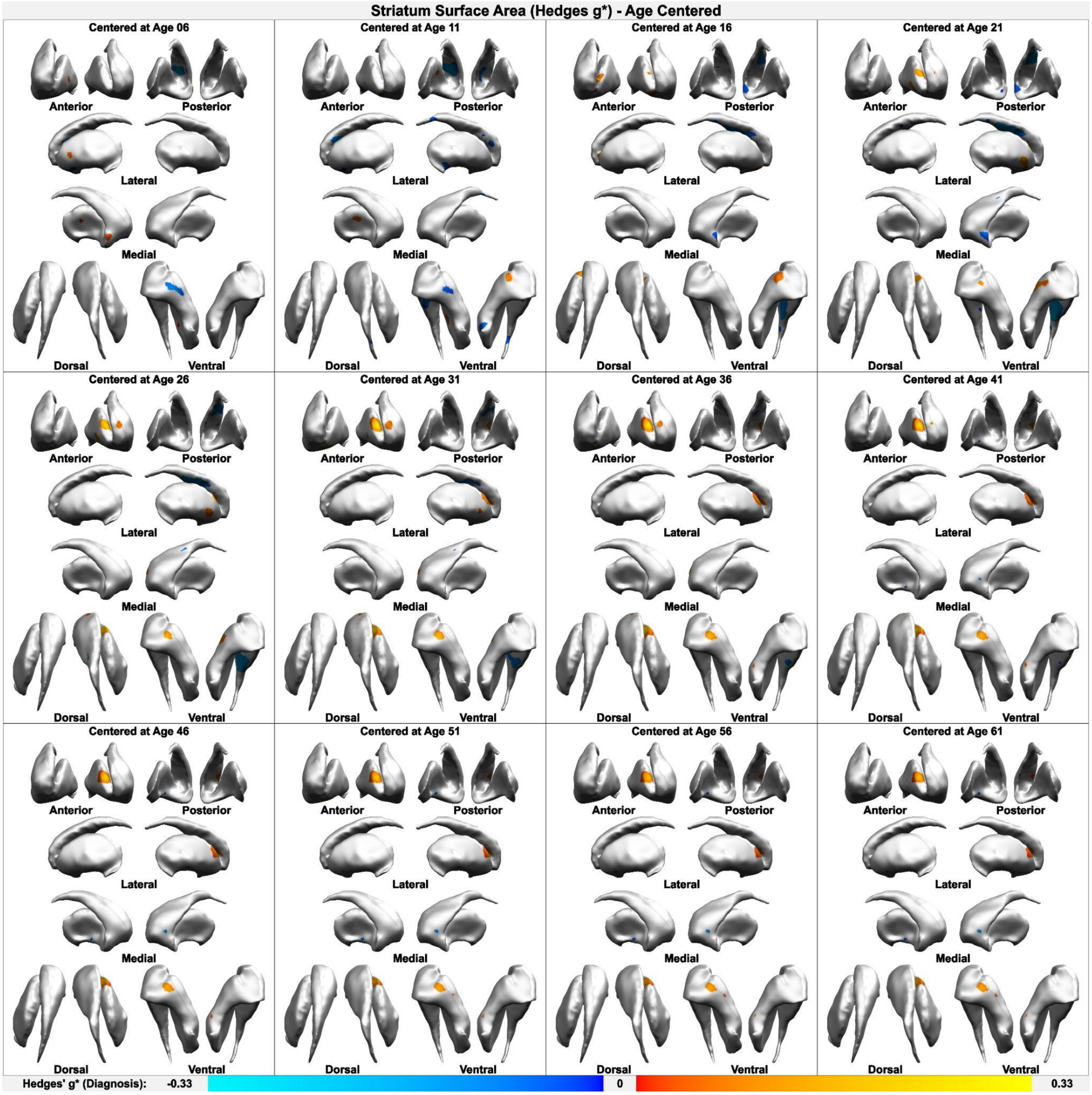


**Supplementary Figure SF-9**: Age-centered analysis, centered on five-year intervals from ages 6-61. Hedges' *g** main effect of ASD diagnosis on vertex-wise surface area in the left and right striatum, when controlling for total structure volume and sex. Warm colours indicate positive effects, cool colours indicate negative effects (*g** range -0.33 to 0.33). Surface area is the Voronoi area surrounding a vertex.

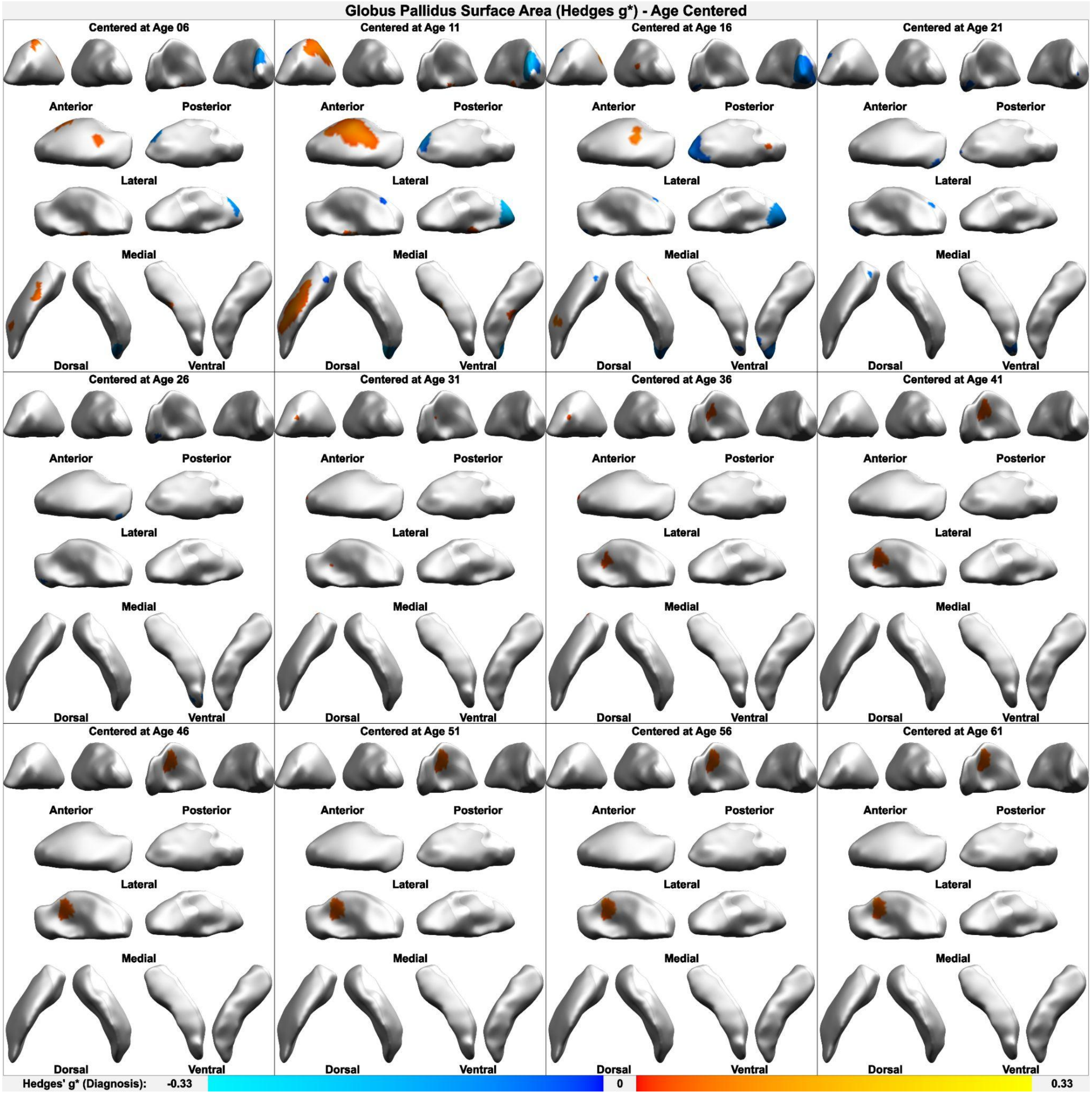


**Supplementary Figure SF-10**: Age-centered analysis, centered on five-year intervals from ages 6-61. Hedges' $g*$ main effect of ASD diagnosis on vertex-wise surface area in the left and right globus pallidus, when controlling for total structure volume and sex. Warm colours indicate positive effects, cool colours indicate negative effects ($g*$ range -0.33 to 0.33). Surface area is the Voronoi area surrounding a vertex.

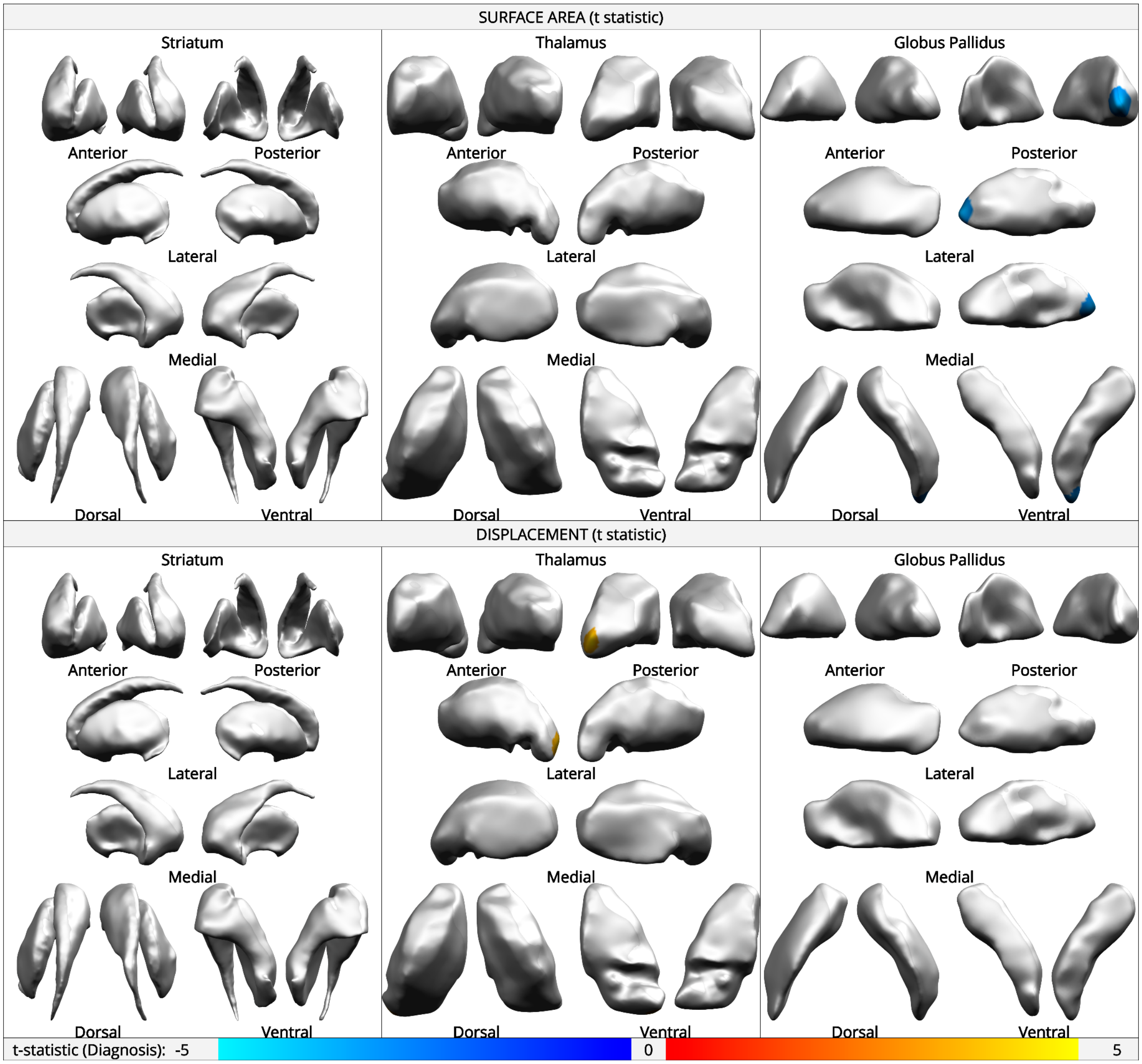


**Supplementary Figure SF-11**: Linear-mixed model mega-analysis. Main effect of ASD diagnosis on vertex-wise surface area and displacement in the left and right striatum, thalamus, and globus pallidus, when controlling for total structure volume, age, and sex. Warm colours indicate positive effects, cool colours indicate negative effects (t-value range -5 to 5). Surface area is the Voronoi area surrounding a vertex; displacement represents relative convexity (positive) or concavity (negative). Thresholded at FDR < .05.

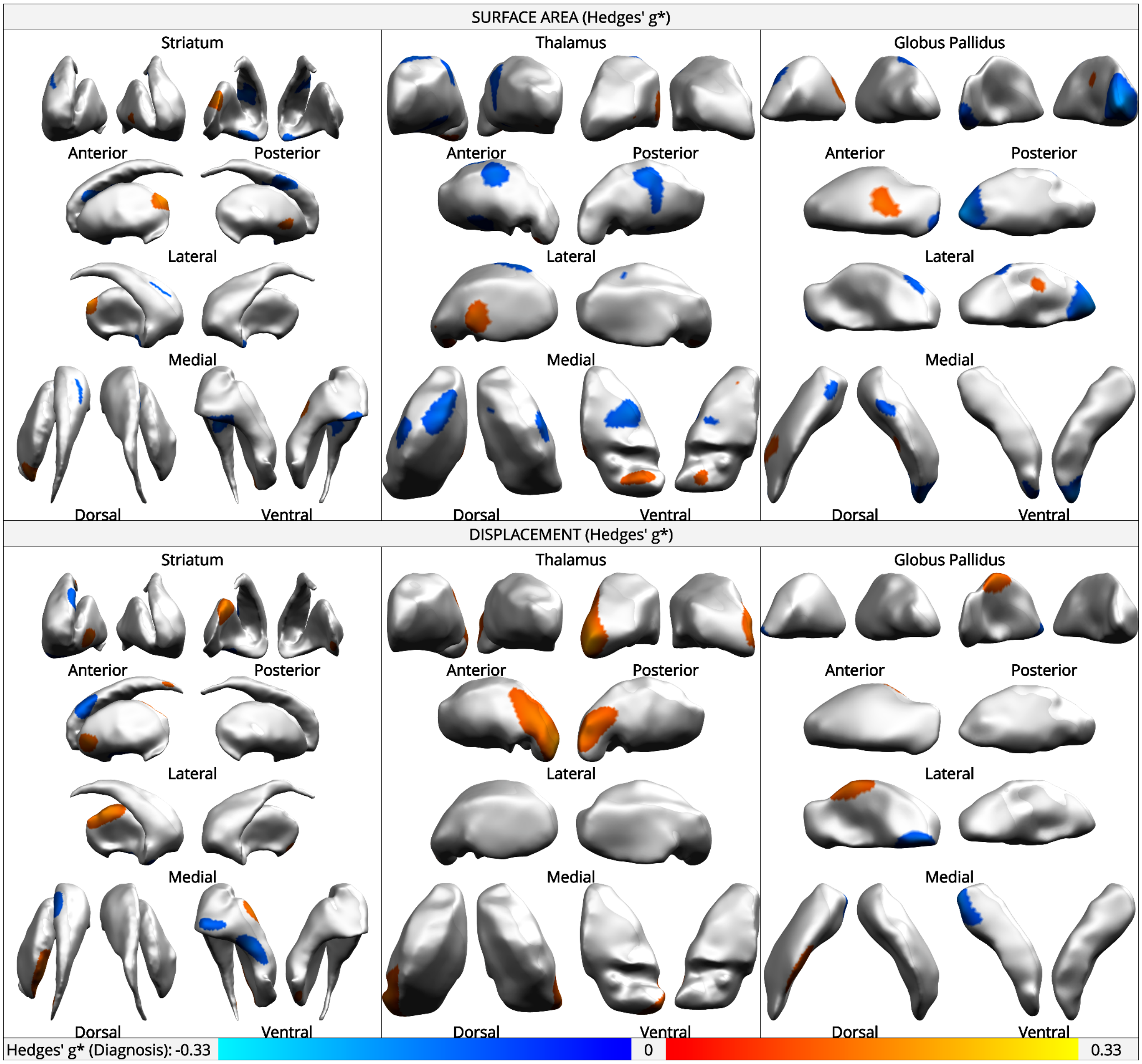


**Supplementary Figure SF-12:** Uncorrected for multiple comparisons, ComBat-harmonized mega-analysis without controlling for FIQ. Main effect of ASD diagnosis, on vertex-wise surface area and displacement in the left and right striatum, thalamus, and globus pallidus, when controlling for total structure volume, age, and sex. Warm colours indicate positive effects, cool colours indicate negative effects (Hedges' g* range -0.3 to 0.3). Surface area is the Voronoi area surrounding a vertex; displacement represents relative convexity (positive) or concavity (negative). Thresholded at $p < .05$. No effects survived FDR multiple comparisons correction.

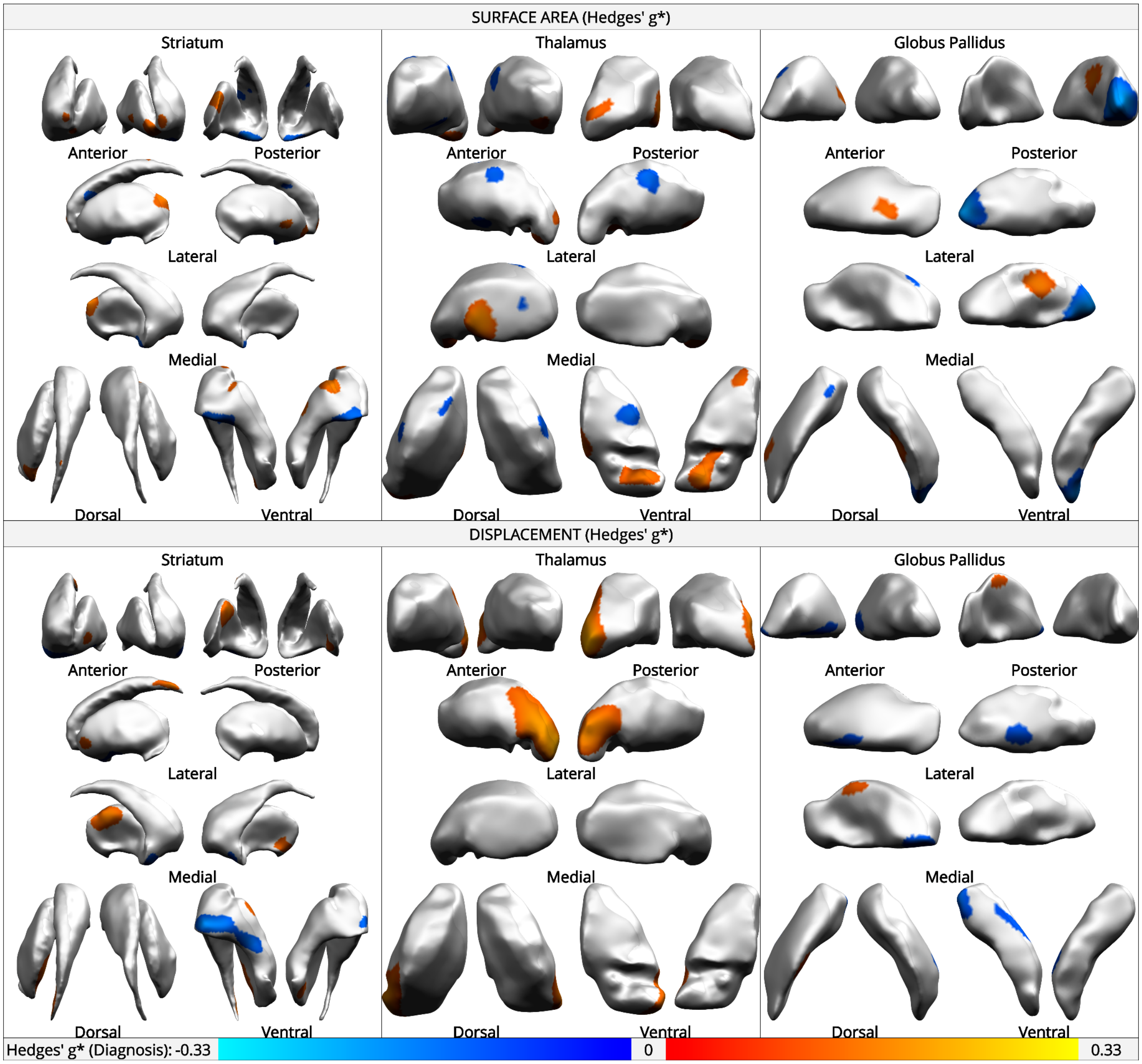


**Supplementary Figure SF-13**: Uncorrected for multiple comparisons, ComBat-harmonized mega-analysis, controlling for FIQ. Main effect of ASD diagnosis, on vertex-wise surface area and displacement in the left and right striatum, thalamus, and globus pallidus, when controlling for total structure volume, age, sex, and FIQ. Warm colours indicate positive effects, cool colours indicate negative effects (Hedges' $g^*$ range -0.3 to 0.3). Surface area is the Voronoi area surrounding a vertex; displacement represents relative convexity (positive) or concavity (negative). Thresholded at $p < .05$. No effects survived FDR multiple comparisons correction.

#### Supplementary Table ST-1

| | Total | Motion Passed | L Str Passed | R Str Passed | L GP Passed | R GP Passed | L Thal Passed | R Thal Passed |
|---|---|---|---|---|---|---|---|---|
| **ABIDE - IP** | 56 | 39 | 26 | 31 | 38 | 38 | 38 | 38 |
| **ABIDE - KKI** | 265 | 156 | 119 | 117 | 156 | 156 | 156 | 156 |
| **ABIDE - MAX MUN** | 57 | 41 | 28 | 34 | 41 | 41 | 37 | 39 |
| **ABIDE - NYU** | 289 | 195 | 148 | 157 | 195 | 195 | 190 | 188 |
| **ABIDE - OHSU** | 121 | 98 | 77 | 76 | 98 | 98 | 96 | 95 |
| **ABIDE - SDSU** | 94 | 48 | 39 | 41 | 47 | 47 | 47 | 47 |
| **ABIDE - UM** | 145 | 66 | 44 | 46 | 66 | 66 | 62 | 63 |
| **CFSA** | 96 | 57 | 39 | 35 | 57 | 57 | 53 | 54 |
| **UKAIMS - Cambridge** | 126 | 114 | 82 | 81 | 114 | 114 | 109 | 110 |
| **UKAIMS - IoP** | 127 | 105 | 68 | 77 | 105 | 105 | 99 | 97 |
| **TORONTO** | 521 | 403 | 254 | 270 | 402 | 387 | 366 | 374 |
| ***Total*** | 1897 | 1322 | 924 | 965 | 1319 | 1304 | 1253 | 1261 |

Segmentation results for original MAGeT Brain workflow, showing the number of participants per site, including total number, number that passed motion QC, and number that passed segmentation QC for each structure. Only sites that were not excluded are shown.
